# The MLL3/TP53/PIK3CA cancer driver mutations promote HIF1α-dependent recruitment and differentiation of pro-tumor ICOS^hi^GITR^hi^ Blimp-1^+^ effector regulatory T cells in breast tumors

**DOI:** 10.1101/2022.10.02.510540

**Authors:** Marie Boutet, Kenta Nishitani, Piril Erler, Nicole Couturier, Zheng Zhang, Anna Maria Militello, Marcelo Coutinho De Miranda, Emeline Barbieux, Erik Guillen, Masako Suzuki, Joseph A. Sparano, Cristina Montagna, Wenjun Guo, Gregoire Lauvau

**Author notes:** **Correspondence to GL or WG: GL:** Albert Einstein College of Medicine, Department of Microbiology and Immunology, 1301 Morris Park Avenue, Bronx, NY, 10461, USA., **WG:** Albert Einstein College of Medicine, Department of Cell Biology, Ruth L. and David S. Gottesman Institute for Stem Cell and Regenerative Medicine Research, 1301 Morris Park Avenue, Bronx, NY 10461, USA. These authors contributed equally (Marie Boutet, Kenta Nishitani, Piril Erler, Wenjun Guo, Gregoire Lauvau). **One Sentence Summary:** MLL3 loss promotes tumorigenesis by enhancing recruitment of effector regulatory T cells.

## Abstract

While essential gatekeepers of immune homeostasis, Foxp3^+^ regulatory T (T_reg_) cells infiltrating tumors acquire distinct phenotypes and become highly immunosuppressive, promoting tumor immune escape and growth. How this occurs and relates to tumor-driver mutations is largely uncharacterized. Herein, we created a mouse mammary stem cell-based tumor model using CRISPR gene editing in which we introduced known human cancer-driver mutations. These included functional loss of the MLL3 histone methyltransferase and p53, and constitutive PI3-kinase activation, recapitulating the genetic makeup of aggressive breast cancers. We show that MLL3 loss fosters tumorigenesis by promoting the rapid establishment of an immunosuppressive microenvironment through induction of HIF1α, which increases the secretion of the chemokine CCL2 by tumor cells and the recruitment of higher numbers of Foxp3^+^ T_reg_ cells via CCR2. Greater infiltration of T_reg_ cells also correlates with MLL3 downregulation and mutations in human breast cancer biopsies. Interestingly, HIF1α enforces the differentiation of tumor-infiltrating T_reg_ cells into highly immunosuppressive ICOS^hi^GITR^hi^ Blimp-1^hi^ effector T_reg_ cells that enable rapid tumor escape. Monoclonal antibody targeting of ICOS or GITR inhibits tumorigenesis in most mice even two months after the cessation of treatment as well as the growth of established tumors, suggesting possible therapeutic opportunities for MLL3-mutant breast cancers.

## Introduction

The relationship between tumor cells and the immune system is complex and involves multiple mechanisms. The concept of cancer immunoediting has been proposed as a framework to account for the processes resulting in the formation of established tumors and subsequent cancer progression^1,2^. Perhaps the most critical step in this model is when tumors have accumulated enough cancer driver mutations to breach the equilibrium and escape the immune response. Cancer driver mutations provide escaping tumor cells with intrinsic features that enhance their proliferative, migratory and survival potentials. However, how specific tumor mutations alter the immune response to promote faster tumor onset and growth is still poorly understood and an area of active investigations. Only relatively few direct connections between cancer driver mutations in cancer cells and their modulation of the immune response have been reported ^3^. Some of these mechanisms link known cancer driver mutations (e.g., *TP53*, *MYC*, β-catenin pathway) and i) the upregulation of cell-surface checkpoint inhibitory receptors (PD-L1, PD-L2, etc) and innate immune regulators^4,5^, ii) the modulation of MHC class I expression and NK-cell target ligands^6,7^, iii) the increased secretion of chemokines and cytokines involved in the recruitment and differentiation of immune cells^8–10^ such as myeloid-derived suppressor cells (MDSCs), type-2-polarized tumor associated macrophages or Foxp3^+^ regulatory T (T_reg_) cells.

T_reg_ cells, which represent essential guardians of immune homeostasis, however, facilitate tumor cell escape and the establishment of robust immunosuppression, and their abundance is usually associated with tumor progression^11,12^. Many mechanisms have been reported to explain how T_reg_ cells infiltrate the TME and suppress anti-tumor effector cell responses. The production of chemokines (CCL17/22, CCL1, CCL5, CCL28, CCL2) by tumors cells and other tumor-associated immune or stromal cells contribute to the selective recruitment of T_reg_ cells expressing cognate chemokine receptors (CCR4, CCR8, CCR5, CCR10, CCR2) across different cancers (e.g., lymphomas, hepatocarcinoma, lung, breast and ovarian cancers)^13–18^. Once in the tumors, T_reg_ cells often expand in response to cognate Ag and acquire highly immunosuppressive features, although underlying mechanisms are not well defined. T_reg_ cells have a lower threshold of Ag-mediated activation^19^ and, in the tumor, exhibit metabolic readiness to turn on oxidative phosphorylation and use lactate to support metabolic demands^20–23^. Acquisition of more suppressive features is characterized by high cell-surface expression of inhibitory receptors (e.g., PD-1, CTLA-4 and others)^24–26^, the high affinity IL-2 receptor alpha chain CD25, secretion of immunosuppressive cytokines (IL-10, TGFβ, IL-35) and vEGF^27^ contributing to tumor angiogenesis – all pathways representing potentially selective therapeutic targets for tumor infiltrating T_reg_ cells. Abundant metabolites and hypoxia in the TME can drive T_reg_ cell expansion and differentiation, yet how exactly selective tumor driver mutations, alone or in combination, control these processes is largely unknown. These mechanisms differ depending on the mutational profile of the tumor; therefore it is paramount to achieve systematic understanding of these processes to design novel precision medicine strategies to eradicate distinct subtypes of tumors with various sets of cancer driver mutations.

Breast cancers remain the leading cause of cancer death in women despite significant improvement in early detection and treatment^28^. The five year survival rate for advanced breast cancers is only ∼29%^29^. Recently, immune checkpoint blockade (ICB) therapy has been approved for triple-negative breast cancer treatment, but only a small fraction of breast cancer patients responds to such therapy. Thus, a deeper understanding of the molecular determinants promoting tumor development and enabling suppression of anti-tumor immune response remains a necessity in breast cancer. The *MLL3* gene encoding a histone methyltransferase (also known as *KMT2C*) is among the most frequently mutated genes in various human cancer types^30–32^. Interestingly, the majority of *MLL3* mutations in breast cancer are truncating point mutations or gene deletions that cause loss of function^33–35^. Up to 27% of human breast cancers harbor loss-of-function mutations of MLL3 which correlate with poor prognosis in patients, but there is no clear bias towards specific breast cancer subtypes. These data suggest that MLL3 is a tumor suppressor affecting a large percentage of breast cancers of all subtypes. Functional studies in mouse models and human cancer cell lines showed that the loss of MLL3 promotes tumor onset and growth in several cancer types, including breast, urethral and leukemia ^36–39^. However, the mechanisms by which MLL3 suppresses tumor onset remain poorly defined. To date, MLL3 has been implicated in cell type-specific gene expression during differentiation, such as adipogenesis, myogenesis and macrophage activation^40^. In breast cancer cell lines, MLL3 has been reported to regulate estrogen receptor (ER)-mediated transcription^41,42^. Yet, the high mutation rate of MLL3 in ER-negative breast cancer subtypes suggests that MLL3 controls tumorigenesis through additional mechanisms other than cell-intrinsic lineage differentiation.

In this study, we created a preclinical mouse model of breast cancer by genetically engineering mammary stem cells (MaSCs) to harbor the frequently associated cancer driver mutations *Pi3kca, Trp53^KO^* (referred to as P5) or *Pi3kca, Trp53^KO^* and *Mll3* loss-of-function (referred to as MP5). The modified MaSCs were next implanted in mammary fat pads to investigate the role of MLL3 in tumorigenesis. We report that while the loss of MLL3 promotes faster tumor onset and growth via enhanced tumor cell-intrinsic proliferative capacity, it also promoted the infiltration of a higher number of Foxp3^+^ T_reg_ cells and their differentiation into ICOS^hi^GITR^hi^ Blimp-1^+^ effector T_reg_ cells that accelerated tumor initiation. Mechanistically, MLL3 loss activated HIF1α which enhanced i) transcription of the chemokine CCL2 accelerating T_reg_ cell tumor infiltration and ii) T_reg_ cell-differentiation into ICOS^hi^GITR^hi^ effector T_reg_ cells in the tumor microenvironment (TME). Interestingly, and as a direct link to these findings, the therapeutic targeting of ICOS and GITR inhibited >60% of Mll3-mutant tumor onset and growth. Collectively, these results represent a first step towards understanding how MLL3, together with p53 loss and PIK3CA activation, regulate the immune microenvironment in breast cancer, and likely that of other cancers where functional MLL3 is lost.

## RESULTS

### Loss of *Mll3* cooperates with Pik3ca activation and *Trp53* deletion to drive tumor initiation and growth

We previously established that *Mll3* deletion alone is insufficient to initiate tumor formation^39^, suggesting the need for combining this mutation with other clinically relevant cooperating mutations to study *Mll3* loss in breast cancer. Thus, to define the most frequent mutations associated with *MLL3*, we first analyzed patient data from the MSKCC-IMPACT Breast Cancer database containing 1,237 advanced stage patients ^43^. This revealed that *PIK3CA* mutations were the most frequent co-occurring mutations in the *MLL3*-mutant tumors (47.3% of the cases). Furthermore, tumors with both *MLL3* and *PIK3CA* mutations most often harbored *TP53* mutations (34% of the cases) (**Fig. 1a**). The triple-mutant tumors belonged to heterogenous subtypes of breast cancers, similar to the general breast cancer population^44^, including Her2+ (24.5%), triple-negative (22.2%), and estrogen receptor positive (ER+) (54.3%), suggesting a non-subtype-restricted role of *MLL3* mutations in breast cancer **(Suppl. Fig. 1a).** Furthermore, patients with all three mutations had significantly shorter overall survival than those with only *PIK3CA* and *TP53* mutations or without any of these mutations (**Fig. 1b**). Altogether, these patient data suggested that *PIK3CA* and *TP53* mutations are clinically relevant cooperating mutations in *MLL3*-mutant breast cancers.

**Figure 1.**
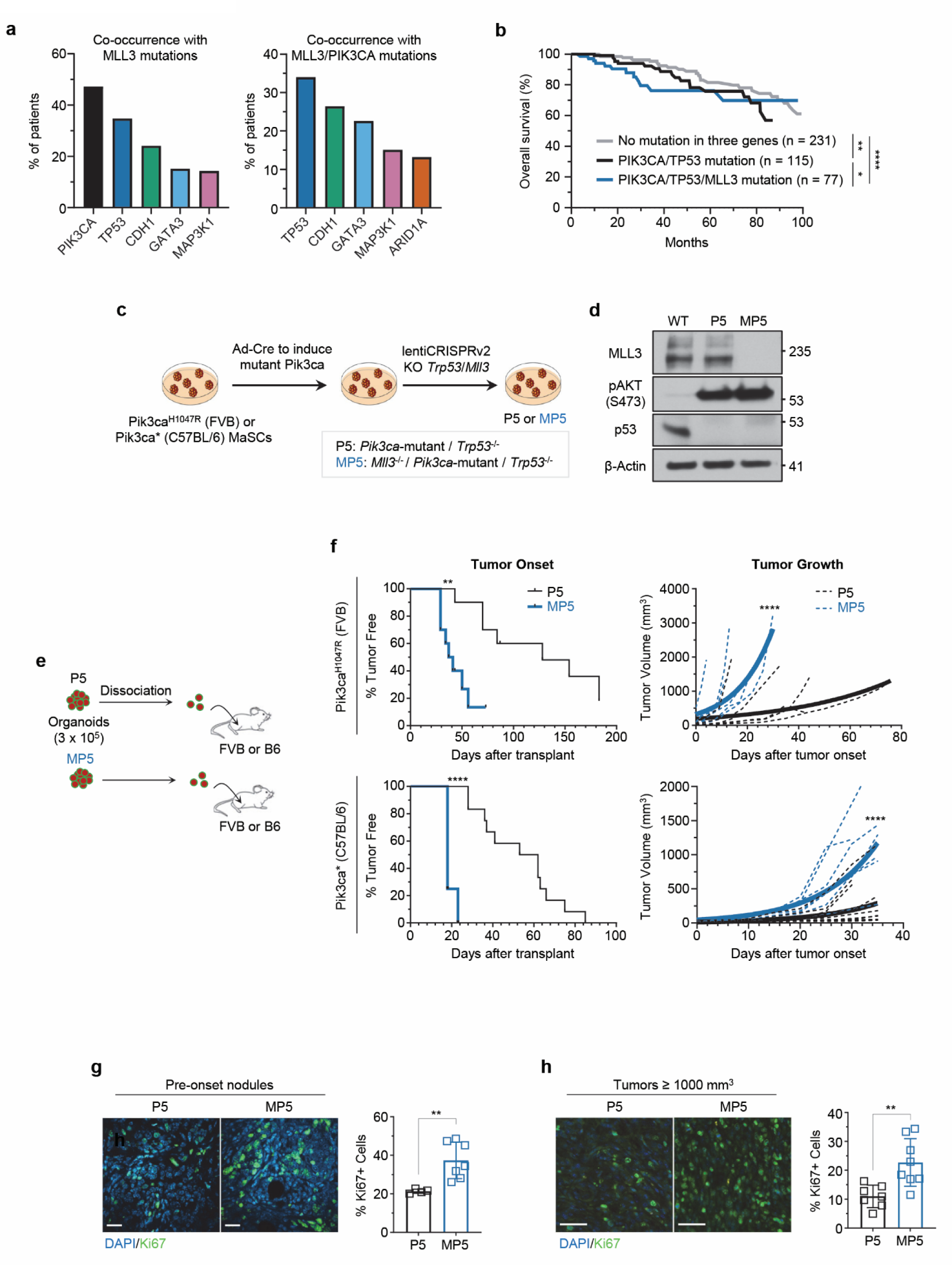
Loss of MLL3 cooperates with PIK3CA and p53 mutations to drive tumor escape and growth. **(a)** Common mutations co-occurring with *MLL3* mutations (left) and with both *MLL3*/*PIK3CA* mutations (right) in patients in MSKCC-IMPACT 2017 Breast Cancer database. **(b)** Overall survival of patients with the indicated mutations in the TCGA Pan Cancer Atlas Breast Cancer dataset. **(c)** Generation of *Pik3ca*-mutant/*p53*^-/-^ (P5) and *Pik3ca*-mutant/*p53*^-/-^/*Mll3*^-/-^ (MP5) mouse mammary stem cell (MaSC) organoids. **(d)** Western blot validation of Pik3ca activation mutation, MLL3 deletion, and p53 deletion in MaSC organoids. **(e)** Schematic of experiment: single cell MaSC were transplanted to mammary fat pads of recipient FVB or B6 WT mice. **(f)** Kaplan-Meier survival analysis of tumor onset (left), and tumor growth rate analysis (right) for P5 tumors (n=10) compared to MP5 tumors (n=10) expressing *Pik3ca*^H1047R^ (line #1, representative of 3 biological replicates) (top) and P5 tumors (n=12) compared to MP5 tumors (n=8) expressing *Pik3ca*\* (line #7, representative of 2 biological replicates) (bottom). **(g)** Immunofluorescence images and quantification of Ki67 expression in P5 pre-onset tumor nodules (n=4) compared to matched MP5 pre-onset tumor nodules (n=7) from line #3. **(h)** Immunofluorescence images and quantification of Ki67 expression in P5 tumors (n=7) compared to MP5 tumors (n=8) (combined data from lines #1 and #3; tumors larger than 1000mm^3^). Data are presented as mean ± SEM. P-values were determined by Log-rank test (b and f) or unpaired t-test (g and h). **p< 0.01, ****p< 0.0001, ns or not indicated: non-significant.

Next, we used our previously established mammary stem cell (MaSC)-based genetically engineered mouse model (GEMM) approach^39^ to generate MaSC-GEMMs carrying either only *Pik3ca* and *Trp53* mutations (P5) or all three mutations combined (MP5) with the goal to study the effect of MLL3 loss during tumorigenesis in a clinically relevant genetic setting. Briefly, we isolated MaSCs from mice that carry a Cre-inducible constitutively active patient hotspot mutation *Pik3ca*^H1047R^ (FVB background)^45^ or a constitutively active mutant *Pik3ca\** (C57BL/6 background)^46^. Next, we activated the mutant *Pik3ca* gene using an adenoviral Cre vector, and sequentially knocked-out the *Trp53* and *Mll3* genes with lentiviral CRISPR (**Fig. 1c**). Both gene deletions and *Pik3ca* gene activation were validated by western blot, showing loss of p53 and MLL3 protein expression and upregulation of phospho-AKT, a downstream target of PIK3CA (**Fig. 1d**). As expected, the MP5 cells showed decreased levels of histone H3 lysine 4 mono-methylation (H3K4me1) and lysine 27 acetylation (H3K27ac) compared to P5 cells, which are known functions of the MLL3 complex^30^ **(Suppl. Fig. 1b, c)**.

We then tested the effect of MLL3 loss on tumor initiation and progression *in vivo* through orthotopic transplantation of P5 and MP5 MaSCs to mammary glands of syngeneic wild-type (WT) recipient mice matching the strain background of MaSCs (**Fig. 1e**). Injection of MP5 MaSCs led to much earlier tumor onset in comparison to P5 cells (**Fig. 1f**). Furthermore, MP5 tumor growth was significantly faster than that of P5 tumors. These results were confirmed across 5 pairs of independently engineered P5 and MP5 MaSC lines that expressed either the *Pik3ca*^H1047R^ or *Pik3ca\** mutation^46^. On average, median tumor onset and volume doubling time were 2.3±0.7 and 2.9±1.1 fold faster in MP5 than P5 tumors, respectively (**Fig. 1f, Suppl. Fig. 1d)**. Consistent with the faster onset and growth of MP5 tumors, the frequency of Ki67^+^ cells in MP5 tumors was significantly higher than in P5 tumors at both pre-onset (1-2 mm diameter) and large-tumor (≥ 1000 mm^3^) stages (**Fig. 1g, h**). In contrast, apoptosis, as measured by cleaved caspase 3, was comparable between P5 and MP5 tumors at both stages **(Suppl. Fig. 1e, f)**. Thus, this model, which recapitulated human breast cancer genetic makeup, establishes that the loss of MLL3, combined with the most frequent co-occurring mutations, loss of *Tp53* and activated *Pik3ca*, enhances tumor onset and growth, which is consistent with the poor overall survival observed in human breast cancer patients with mutations in all three genes.

### Foxp3^+^ regulatory T cells rapidly infiltrate *Mll3*-mutant tumors during mouse tumor initiation and in patients

To explore whether MLL3 loss modulates tumor-infiltrating immune cells during tumor onset, we next characterized the composition of immune cells infiltrating P5 and MP5 tumors at very early stages of tumor formation using high dimensional spectral flow cytometry with a panel of 27 distinct markers we developed and that includes all major immune cell lineages (**Table S1**). WT mice, either on FVB or C57BL/6 (B6) genetic backgrounds, were injected with P5 or MP5 MaSCs, which were allowed to grow for 20 and 10 days, respectively, until they formed comparable sized nodules (1-2 mm) prior onset of palpable tumors (∼3 mm diameter) (**Fig. 2a and Supp. Fig. 2a**). Pre-onset tumor nodules were isolated and dissociated for FACS analysis of immune cell infiltrates. We gated cells based on forward and side scatter, excluded cell doublets and focused on CD45^+^ hematopoietic-derived immune cells that either expressed or not the myeloid marker CD11b (**Fig. 2b**). Next, we used t-Distributed Stochastic Neighbor Embedding (t-SNE) for unsupervised clustering and dimensionality reduction, and the identification of distinct or similar clusters of cells and their relative proportions. T-SNE analysis and population quantification across individual mice revealed that the proportion of pre-onset tumor-infiltrating CD45^+^ leukocytes including lymphocytes (CD4^+^ T, CD8^+^ T, B and NK cells) and subsets of myeloid cells (granulocyte-derived CD11b^+^Ly6G^+^, monocyte-derived CD11b^+^Ly6C^hi^, and Ly6C/G^neg^CX3CR1^+^) were comparable between P5 and MP5 pre-onset tumor nodules in WT FVB and B6 mice (**Fig. 2c and Suppl. Fig. 2a**). Numbers of CD45^+^ cells and all various immune cell subsets were also equivalent in P5 and MP5 nodules. Interestingly, however, we revealed significantly higher proportions and numbers of Foxp3^+^ CD4^+^ regulatory T (T_reg_) cells in MP5 compared to P5 nodules (2-fold, **Fig. 2d and Suppl. Fig. 2a)**. Immunofluorescence (IF) of pre-onset tumor nodule sections also confirmed the significant increase in T_reg_ cells in MP5 compared to P5 nodules (**Fig. 2e**). Of note, T cell subsets in tumor-draining LNs including T_reg_ cells, were comparable, in line with the implication of tumor-intrinsic mechanisms to account for increased tumor infiltrating T_reg_ cells in MP5 versus P5 nodules (**Supp. Fig. 2b**).

**Figure 2.**
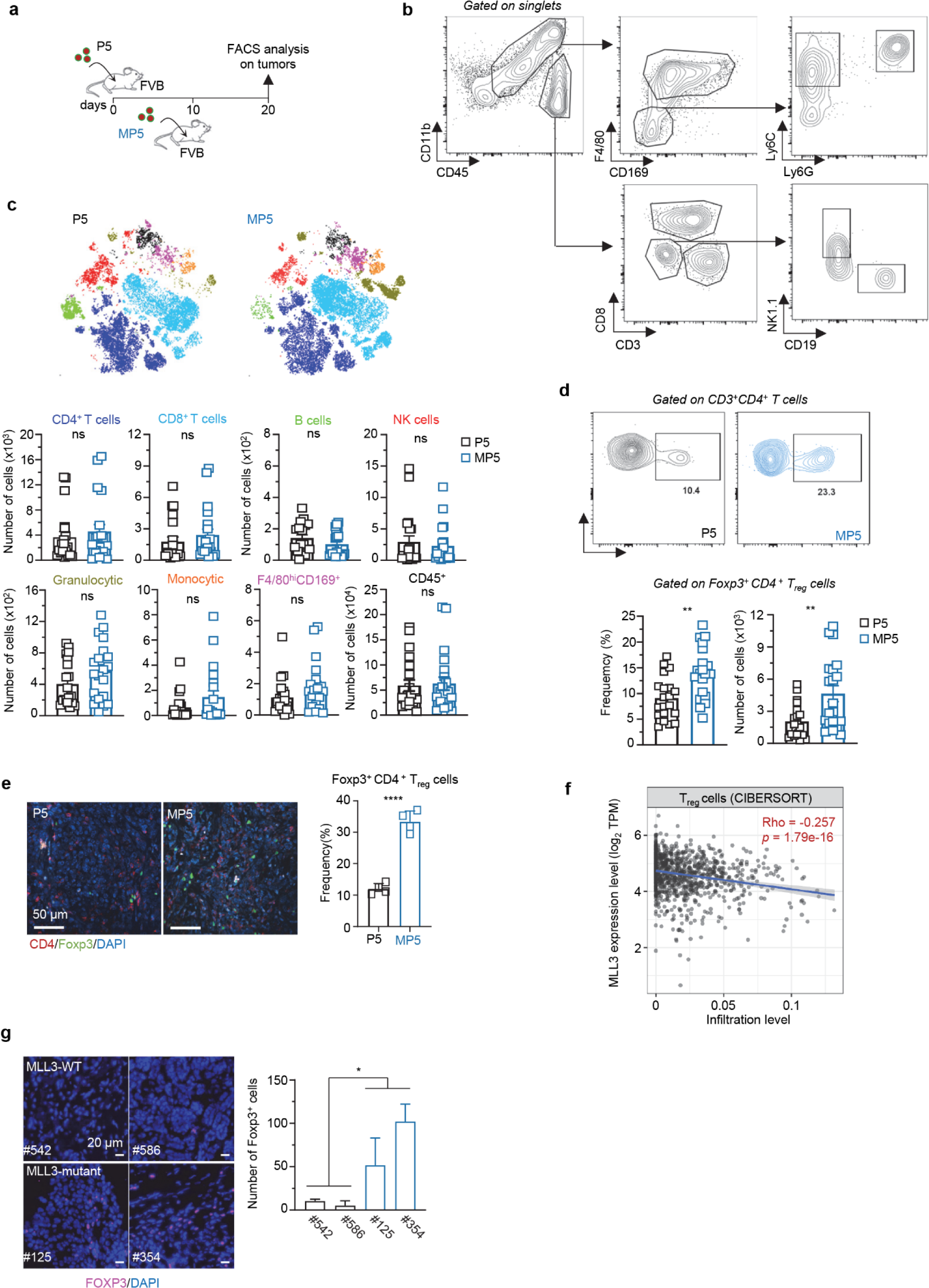
Loss of MLL3 promotes T_reg_ cell infiltration during tumor initiation and correlates with higher T_reg_ cell frequency in patients. **(a)** Schematics of experiment: 3×10^5^ P5 or MP5 MaSCs from line #3 were transplanted into FVB mice. Pre-onset tumor nodules were extracted and analyzed by flow cytometry (20- and 10-days post-transplant, respectively). **(b)** Tumor samples were stained for immune cell lineages and analyzed by flow cytometry. Example of cell-lineage gating strategy to define immune cell subsets. **(c)** T-SNE overlays of subsets of immune cells as defined by the color codes. **(d)** Graphs show the frequency and the number of Foxp3^+^CD4^+^ T_reg_ cells with representative dot plots. Data (c, d) are from 4 independent replicate experiments (P5, n=19 and MP5, n=22 pre-onset tumor nodules from line #3). **(e)** Immunofluorescence images and quantification of CD4 and Foxp3 co-expression in sections of P5 compared to MP5 pre-onset tumor nodules at 20- and 10-days post-transplant, respectively (n=4, P5 and MP5 pre-onset tumor nodules from line #3). **(f)** Correlation between *MLL3* expression (log_2_ TPM) and T_reg_ infiltration level in breast cancer patients by TIMER2.0 analysis (n=1100). **(g)** Immunofluorescence images and quantification of Foxp3^+^ T_reg_ cells performed on fresh frozen breast tumor tissue sections of human breast cancer patients mirroring the TP53, PIK3CA and MLL mutational status of P5 (#542 and #586) and MP5 (#125 and #354). The values were derived from the mean number of 5 distinct fields of view on each slide, and the count was normalized to the total DAPI^+^ area. P-values were determined by unpaired t-test (c,d,e,g, and h) or Spearman’s rank correlation test (f). *p< 0.05, **p< 0.01, ****p< 0.0001, ns or not indicated: non-significant.

Next, we investigated whether T_reg_ cell infiltration was associated with *MLL3* gene expression levels in breast cancer patients. TIMER2.0 (Tumor Immune Estimation Resource) analysis, a bioinformatic tool that quantifies the abundance of immune infiltrates using bulk RNA-seq data^47^, revealed that T_reg_ cell infiltration was negatively correlated to *MLL3* mRNA expression levels in the TCGA samples (**Fig. 2f**), consistent with our findings in the murine model. To further extend these observations, we aimed to identify breast cancer tumor biopsies with concurrent PI3KCA and TRP53 gene mutations with or without mutation in the MLL3 gene. We screened a cohort of 57 patient samples sequenced for cancer driver mutations as we previously described ^48^ and identified two cases containing *MLL3*, *PIK3CA* and *TP53* mutations and two with only *PIK3CA* and *TP53* mutations (**Table S2**). FOXP3 IF on tumor sections of these cases revealed that *MLL3*-mutant breast tumors had substantially more FOXP3^+^ T_reg_ cells than *MLL3*-WT counterparts (**Fig. 2g**). This analysis, despite the limited sample size, supports the finding that MLL3 mutation also contributes to T_reg_ cell infiltration in human breast cancer, consistent with the murine model.

### T_reg_ cells infiltrating the breast tumors exhibit an immunosuppressive effector phenotype compared to those from draining lymph nodes

To further characterize the tumor-infiltrating Foxp3^+^ T_reg_ cells in pre-onset tumors (P5+MP5) and associated draining lymph nodes (dLN), we developed a new Foxp3^+^ T_reg_ cell-focused spectral flow cytometry panel and used subsequent Uniform Manifold Approximation and Projection (UMAP) for dimension reduction analysis (**Table S1**). Both for tumor bearing mice, T_reg_ cells from dLN segregated away from those in tumors in a UMAP analysis (**Fig. 3a**). While dLN T_reg_ cells mostly exhibited a naïve phenotype (CD62L^hi^, CD44^low^) and did not proliferate (Ki67^low^), those infiltrating the tumors were activated, proliferated and co-expressed multiple cell-surface inhibitory receptors such as GITR, ICOS, PD-1, CTLA-4, LAG3 and TIGIT (**Fig. 3b**). Tumor-infiltrating T_reg_ cells also upregulated the TFs blimp-1 and T-bet which drive effector T cell differentiation as well as the integrin CD103 involved in tissue residency. Thus, T_reg_ cells infiltrating the breast tumors acquired an activated effector phenotype that is very distinct from naïve LN T_reg_ cells, and expressed high levels of Blimp-1 and multiple inhibitory receptors, consistent with higher suppressive features.

**Figure 3:**
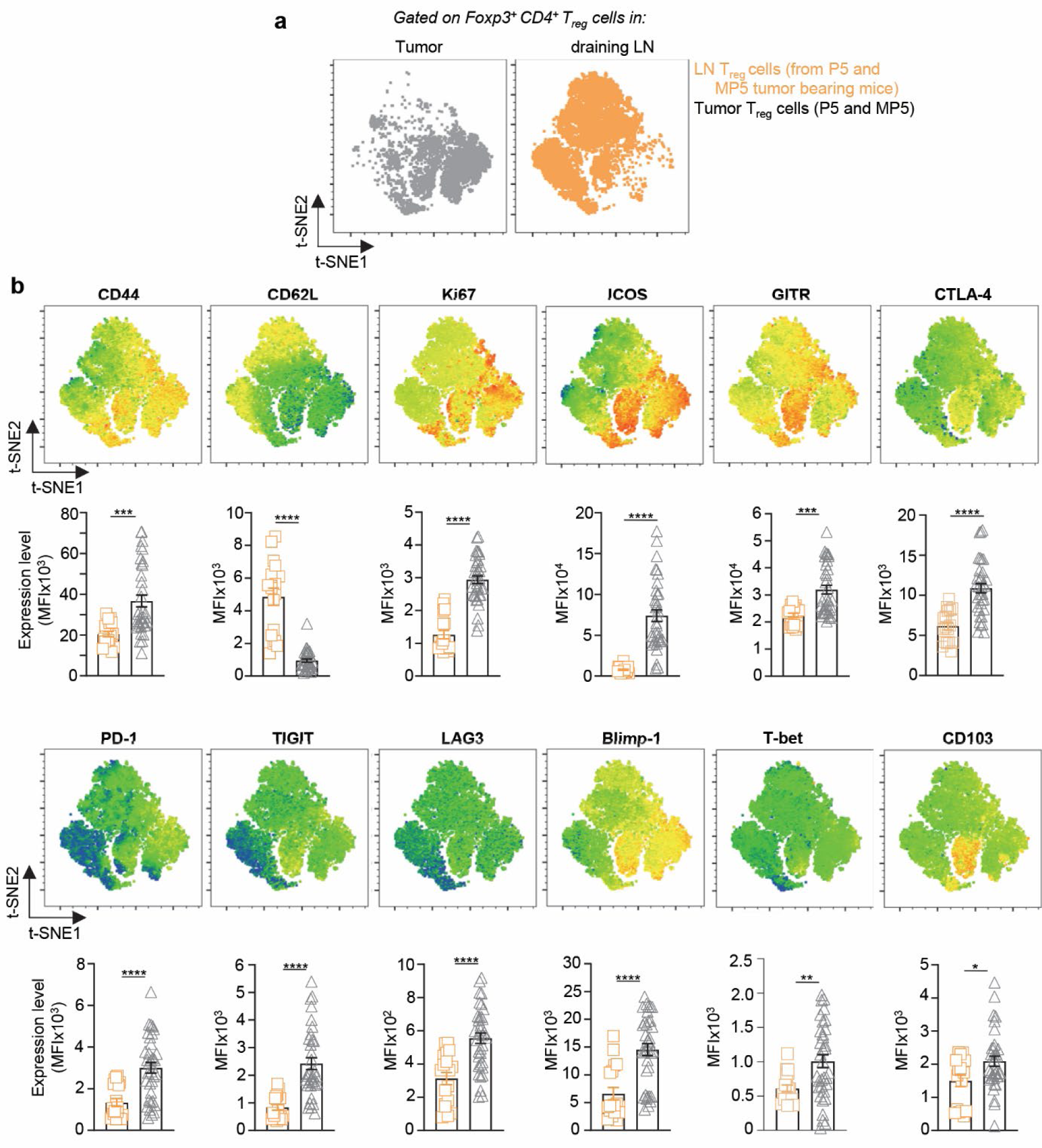
Tumor-infiltrating T_reg_ cells acquire an effector immunosuppressive phenotype. Overlays of t-SNE spatial distribution of T_reg_ cells based on 22-marker flow cytometry analysis in the dLN versus pre-onset tumors of mice transplanted with both P5 and MP5 MaSC. Heat maps of relative expression levels of indicated markers at the single-cell level on corresponding t-SNE maps with bar graphs averaging T_reg_ cells expression level for indicated markers. t-SNE maps concatenate data from 3 independent replicate experiments (P5: n=19 tumors and MP5: n=22 tumors). Each symbol (bar graphs) feature 1 tumor. P-values were determined by unpaired t-test (b). *p< 0.05, **p< 0.01, ***p< 0.001, ****p< 0.0001, ns or not indicated: non-significant.

### Mll3 loss enforces the T_reg_ cell immunosuppressive phenotype and promote faster tumor onset

We next focused the UMAPs on T_reg_ cells from MP5 versus P5 tumors which highlighted further differences in T_reg_ cell distribution and abundance (**Fig. 4a**). Specifically, we noted that in addition to higher numbers, MP5-infiltrating T_reg_ cells included distinct expanded subsets compared to P5-infiltrating T_reg_ cells. While expression of several markers (CD44, CD122, CD27, CCR5, LAG-3, TIGIT, PD-1, T-bet, Eomes) did not differ between P5 and MP5 tumor-infiltrating T_reg_ cells, markers associated with effector cell differentiation (KLRG1, Blimp-1, CD62L), proliferation (Ki67), tissue-residency^50^ (CD103) and improved suppressive T_reg_ cell features (CD25, ICOS, GITR, CTLA-4) were more upregulated in MP5-compared to P5-tumor-infiltrating T_reg_ cells^51–53^ (**Fig. 4b and Supp. Fig. 3a**). We also noted that MP5 tumor-infiltrating CD4^+^ and CD8^+^ T_conv_ cells expressed higher levels of PD-1, ICOS and CTLA-4 than P5 counterparts, making them possible targets of immunosuppression (**Supp. Fig. 3b, c**).

**Figure 4.**
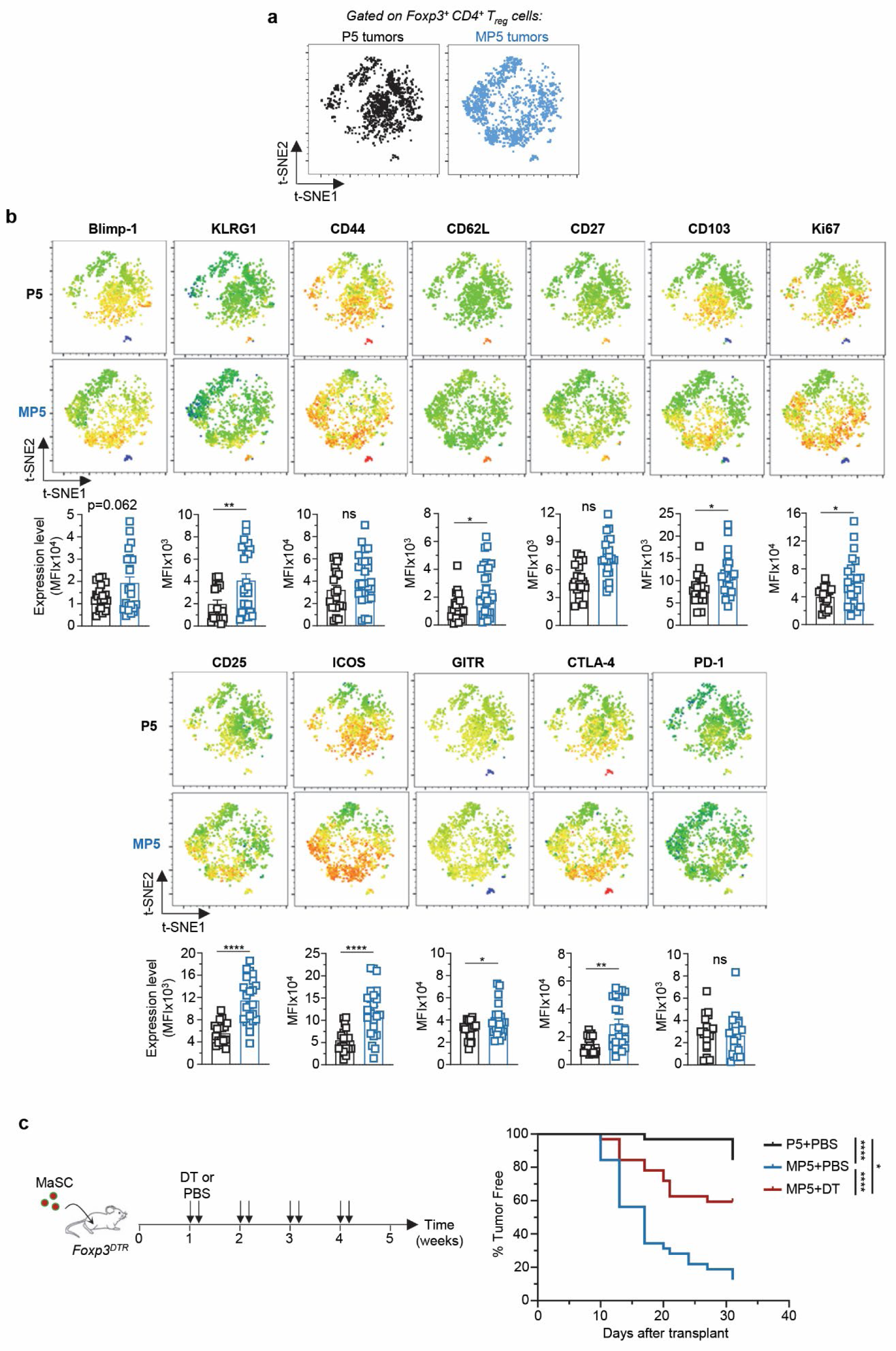
Immunosuppressive T_reg_ cells contribute to faster initiation of Mll3-mutant tumors. **(a, b)** t-SNE spatial distribution of T_reg_ cells infiltrating P5 or MP5 pre-onset tumors based on 22-marker flow cytometry analysis (same data as in Figure 3). **(b)** Relative expression level of indicated markers at the single-cell level on t-SNE maps with bar graphs averaging overall T_reg_ cells expression level for each marker. t-SNE maps concatenate data from 4 independent replicate experiments (P5: n=19 tumors and MP5: n=22 tumors). Each symbol (bar graphs) feature 1 tumor. Schematics of experiment and Kaplan-Meier survival analysis of tumor onset. P5 or MP5 MaSCs (line #2) were transplanted into *Foxp3^DTR^*homozygous mice. Diphtheria toxin (DT) or control PBS injections started one week post MaSC transplant on two consecutive days and then repeated for 3 weeks (n=32 MaSC injections per group, pooled from two independent experiments). P-values were determined by unpaired t-test (b) or Log-rank test (c). *p< 0.05, **p< 0.01, ****p< 0.0001, ns or not indicated: non-significant.

Based on these observations, we hypothesized that T_reg_ cells infiltrating MP5 tumors contributed to establishing faster early immunosuppression and promoted rapid MP5 tumor cell immune escape. We implanted P5 or MP5 MaSCs in *Foxp3^DTR^* mice in which the simian diphtheria toxin receptor (DTR) is knocked-in under the *Foxp3* gene promoter, allowing for selective depletion of Foxp3^+^ T_reg_ cells upon diphtheria toxin (DT) injection. We adopted a previously reported protocol enabling relatively long-term T_reg_ cell depletion ^54^. Briefly, one week after MaSC injection, we administered low-dose DT or PBS twice a week and monitored tumor onset (**Fig. 4c**). We confirmed Foxp3^+^ CD4^+^ T_reg_ cell depletion through flow cytometry analysis of peripheral blood 18 days post MaSC implantation and in the tumors at the endpoint **(Suppl. Fig. 3d)**. T_reg_ cell depletion significantly delayed the onset of MP5 tumors compared to non-depleted mice and to mice implanted with P5 tumors. Although we observed a mild reduction in body weight **(Suppl. Fig. 3e)**, we could follow the mice with T_reg_ cell depletion for at least four weeks after MaSC transplant. Since sustained T_reg_ cell depletion may lead to uncontrolled T_conv_ cell activation and wasting autoimmunity ^55^, we also examined T_conv_ cell activation status by flow cytometry staining of several T cell-surface activation markers (CD11a, CD44, KLRG1, CD62L, CD69, **Suppl. Fig. 3f**). We did not observe any significant upregulation for most of these markers in T_reg_ cell-depleted MP5 tumors, most likely ruling out that the delayed MP5 tumor onset in T_reg_ cell-depleted mice is accounted for by overactivated T_conv_ cells in the TME. Thus, altogether these results suggest that MLL3 loss in breast tumors enhances the differentiation of highly immunosuppressive T_reg_ cells that promote rapid tumor cell escape.

### MLL3 loss induces HIF1α which enhances T_reg_ cell infiltration and tumor onset

Since we had previously established that the HIF1α pathway is the top upregulated pathway induced upon *Mll3* deletion in WT MaSCs ^39^, we hypothesized that HIF1α controls the increased infiltration of T_reg_ cells. Using qRT-PCR analysis, we first confirmed, in multiple independent MaSCs lines, that several HIF1α target genes, such as *Vegfa*, *Glut-1*, *CaIX*, *Timp1* and *Adm* ^56,57^, were also expressed at higher levels in MP5 compared to P5 MaSCs (3-10 fold), ruling out that *Pik3ca* and *Trp53* mutations in MaSCs interfered with the induction of HIF1α subsequent to the loss of Mll3 (**Fig. 5a, Suppl. Fig. 4a**). In addition, immunohistochemistry analysis of pre-onset tumor nodules revealed a marked upregulation of HIF1α expression in MP5 nodules compared to P5 nodules of the same size, and across two pairs of independently engineered MaSCs lines (**Fig. 5b and Suppl. Fig. 4b)**. Analysis of the TCGA Breast Cancer dataset for well-characterized HIF1α target genes further showed that majority of these genes were significantly upregulated in tumors with *MLL3* mutations (**Suppl. Fig. 4c**). Furthermore, TCGA breast tumors with mutations in *PIK3CA*, *TP53*, and *MLL3* genes (MP5-like) exhibited upregulation of the hypoxia gene signature compared to P5-like tumors lacking *MLL3* mutation, consistent with the results in the mouse models (**Fig. 5c**).

**Figure 5.**
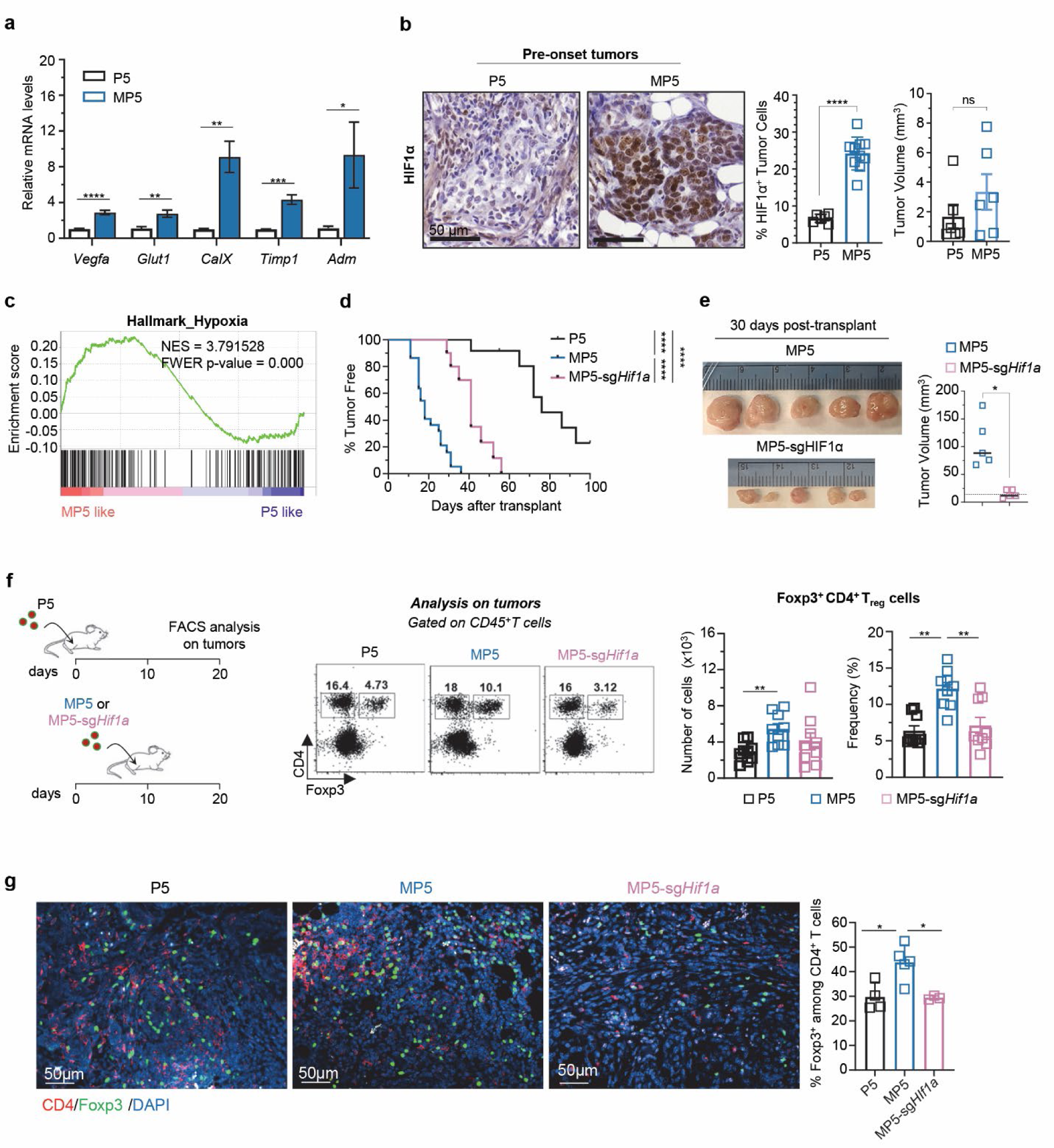
HIF-1α upregulation in *Mll3*-mutant tumors is required for early tumor onset and T_reg_ cell infiltration. **(a)** mRNA expression of HIF1α downstream targets in P5 and MP5 organoids from line #4. **(b)** Immunohistochemistry images and quantification of HIF1α expression in P5 pre-onset tumor nodules (n=6) compared to MP5 pre-onset tumor nodules (n=10) from line #1. Right graph shows tumor volumes of P5 and MP5 tumors analyzed for HIF1α expression. **(c)** Gene Set Enrichment Analysis for Hallmark Hypoxia gene signature in P5- or MP5-like human breast cancer patients from TCGA Pan Cancer Atlas Breast Cancer database (P5-like = 115 patients, MP5-like = 77 patients). **(d)** Kaplan-Meier survival analysis of tumor onset for P5 control (P5-sg*NT*, n=12), MP5 control (MP5-sg*NT*, n=22) or MP5 HIF1α-knockout (MP5-sg*Hif1a*, n=10) MaSCs transplanted to WT FVB mice. Pooled from two independent experiments. **(e)** Dissected tumor images and mean volumes of MP5-sg*NT* (n=5) and MP5-sg*Hif1a* (n=5) tumors collected 30 days post-transplantation. A centimeter ruler scale (with 1mm units) is included in the images. **(f)** Schematics of transplant of P5-sg*NT*, MP5-sg*NT* and MP5-sg*Hif1a* MaSCs in FVB mice with read out outcome. Representative dot plots gated on CD4^+^ T_conv_ cells and Foxp3^+^ T_reg_ cells in pre-onset tumor nodules. Bar graphs show numbers and frequency pooled from 3 independent replicate experiment (n=9). **(g)** CD4 and Foxp3 immunofluorescence images and frequency of Foxp3^+^ cells among CD4^+^ T cells in P5-sg*NT* (n=4), MP5-sg*NT* (n=5) and MP5-sg*Hif1α* (n=3) pre-onset tumor nodules. P-values were determined by unpaired t-test (a,b,e,f, and g) or Log-rank test (d). *p< 0.05, **p< 0.01, ***p< 0.001, ****p< 0.0001, ns or not indicated: non-significant.

Next, to directly test if the upregulation of the HIF1α pathway contributed to the accelerated onset of MP5 tumors and the infiltration of highly suppressive T_reg_ cells, we deleted *Hif1a* in MP5 MaSCs using lentiviral CRISPR, and validated the deletion by western blot (**Suppl. Fig. 4d**). We transplanted *Hif1a*-deleted MP5 (MP5-sg*Hif1a*) and control MP5 and P5 cells into WT recipient mice (**Fig. 5d and Suppl. Fig. 4f**). The onset of MP5-sg*Hif1a* MaSCs was significantly delayed (by ∼23 days) and tumors were markedly smaller (∼7 fold) compared to MP5 MaSCs 30 days post-transplantation (**Fig. 5e**). We note that MP5-sg*Hif1a* MaSCs still exhibited faster tumor onset phenotypes than P5 control, which is likely due to incomplete *Hif1a* deletion as shown in MaSC organoids or pre-onset tumors (**Suppl. Fig. 4d and 4e)**. Importantly, the analysis of immune cell infiltrates 20 (P5) and 10 (MP5 and MP5-sg*Hif1a*) days later, when the three types of cells formed similar sized pre-onset nodules, revealed lower frequencies (∼3 fold less) and numbers (∼2 fold less) of T_reg_ cells in MP5-sg*Hif1a* and P5 nodules compared to MP5 (**Fig. 5f**). This result was also confirmed by immunofluorescence on sections of pre-onset tumor nodules, showing significantly fewer Foxp3^+^ CD4^+^ T_reg_ cells in MP5 tumors lacking HIF1α (MP5-sg*Hif1a*) compared to control tumors (MP5), and proportions were also comparable to P5 tumors (**Fig. 5g**). Moreover, while the overall numbers of tumor-infiltrating CD45^+^ immune cells, CD4^+^ T_conv_ cells, NK cells and Ly6C^+^monocytic cells were comparable in all pre-onset tumors (**Supp. Fig. 4g**), that of CD8^+^ T cells, Ly6G^+^ granulocytic cells and F4/80^+^ MPs was higher in MP5-sg*Hif1a* tumor nodules, indicating that loss of HIF1α more specifically alters T_reg_ cell infiltration compared to other CD45^+^ leukocytes. Thus collectively, these results indicated that *Mll3* loss promotes tumorigenesis (measured as faster tumor onset) at least in part through the upregulation of HIF1α and the increased infiltration of T_reg_ cells in pre-onset tumor nodules.

### HIF1α induces CCL2 production by the tumor cells, promoting CCR2^+^ T_reg_ cell recruitment and faster tumor onset

We hypothesized that cytokines and/or chemokines in *Mll3*-mutant TME promoted rapid T_reg_ cell recruitment in pre-onset tumors, establishing robust immunosuppression early on. Using a multiplex bead-based array, we screened a panel of 26 cytokines/chemokines commonly involved in immune-cell function and tumor growth, and quantified their amounts in P5, MP5, and MP5-sg*Hif1a* pre-onset tumor nodules (**Supp. Fig. 5a**). While the levels of several cytokines and chemokines were significantly different between the P5 and MP5 nodules, CCL2 was the only one that also substantially decreased in HIF1α-deficient MP5 tumors (**Fig. 6a**). Similarly, *Ccl2* mRNA levels were consistently lower in P5 compared to MP5 MaSCs, for all independent cell lines generated (**Supp. Fig. 5b**). MP5-sg*Hif1a* MaSCs had also comparable *Ccl2* mRNA levels to P5 MaSCs, and significantly lower than MP5 MaSCs (**Fig. 6b**). Thus, Mll3-deficient cells upregulated *Ccl2* mRNA and chemokine expression in a HIF1α-dependent manner. Consistent with these findings, we found that higher *CCL2* mRNA expression is elevated in *MLL3-mutant* breast cancers in the TCGA patient dataset and correlated with that of *HIF1A* (**Supp. Fig. 5c, d**). Since HIF1α promoted CCL2 expression, we next tested if HIF1α bound to regulatory regions of the *Ccl2* gene. We searched transcription factor binding motifs around the *Ccl2* gene region using the CiiiDER analysis and identified a HIF1β binding site within the first exon of *Ccl2*. Since HIF1β forms a heterodimer with HIF1α to activate target genes, this suggests HIF1α may bind to the same site. We also identified hypoxia response element (HRE) motifs near both the mouse and human *CCL2* gene transcription start sites (**Supp. Fig. 5e)**. In addition, we found three additional putative enhancer regions based on ENCODE annotation and two reported regulatory regions up- or downstream of the *Ccl2* transcription start site (TSS) ^58,59^ (**Supp. Fig. 5f)**. We next performed HIF1α ChIP-qPCR in MP5 cells to determine whether HIF1α bound to any of these putative regulatory regions (**Fig. 6c**). Compared to gene-desert regions used as negative controls, HIF1α was significantly enriched within three adjacent regions near the *Ccl2* TSS including the HIF1β binding site, indicating that HIF1α could act as a direct transcriptional regulator for *Ccl2* expression.

**Figure 6.**
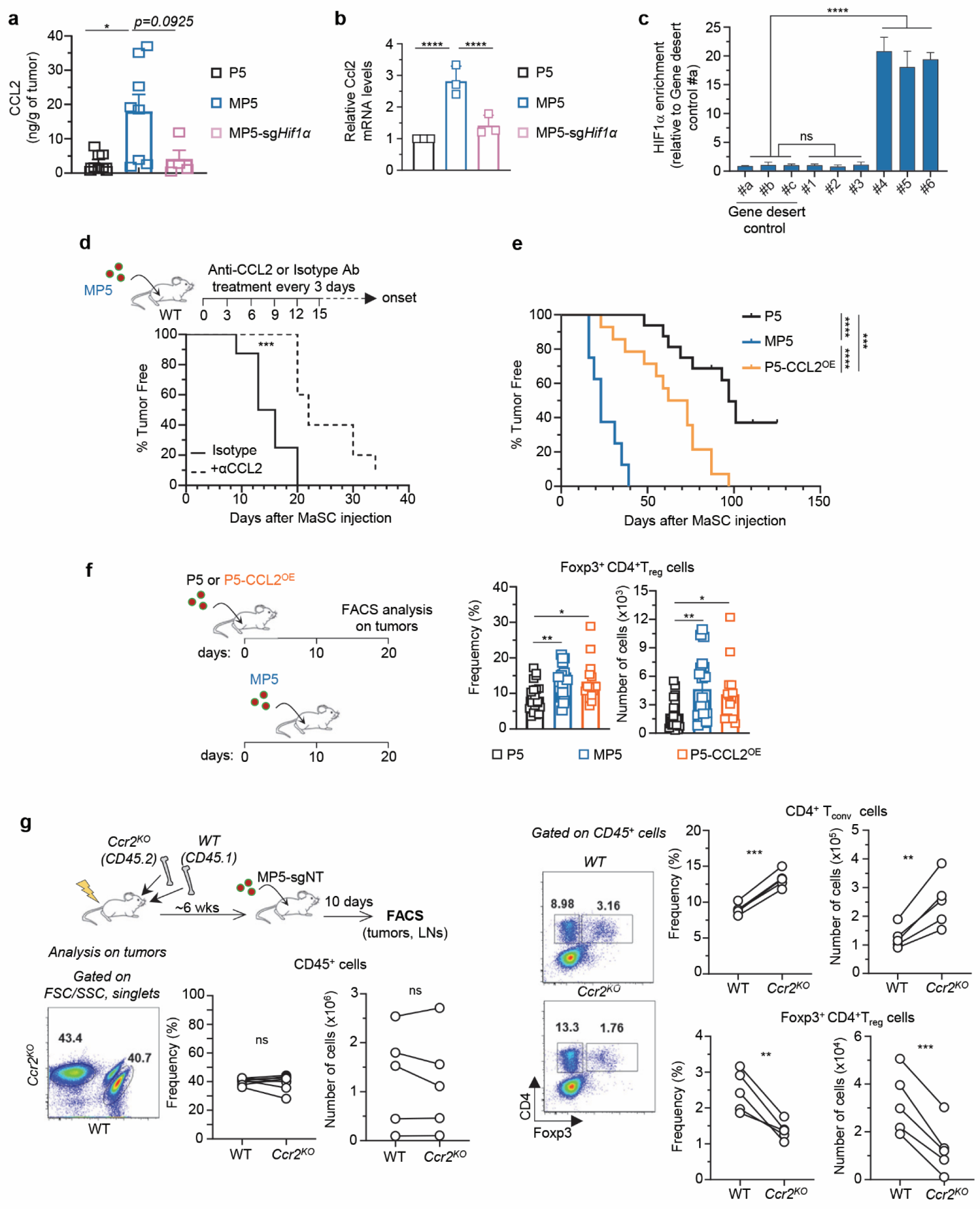
HIF1a mediates upregulation of CCL2 in tumor cells and CCL2 promotes T_reg_ cell infiltration and early tumor onset. **(a)** Quantity of CCL2 chemokine in P5-sg*NT*, MP5-sg*NT*, and MP5-sg*HIF1α* pre-onset tumor nodules. **(b)** *Ccl2* mRNA expression in P5-sg*NT*, MP5-sg*NT* and MP5-sg*Hif1a* organoid lines (line #3). **(c)** Relative enrichment of HIF1α binding around the *Ccl2* transcription start site (TSS) in MP5 MaSCs (line #3). Gene desert regions were used as negative controls. Data are representative from three independent experiments. **(d)** Schematics of MP5 MaSC transplant and mAb treatment strategy in WT B6 mice, and Kaplan-Meier survival analysis of tumor onset for MP5 tumors treated with anti-CCL2 or control isotype mAbs in a pool of 2 experiments (n=10 tumors/group). **(e)** Kaplan-Meier analysis of tumor onset for P5-CCL2^OE^ (n=14) and P5-control vector (n=16) and MP5-control vector (n=8) MaSCs implanted in WT FVB mice. Data are pooled from two independent experiments. **(f)** Schematics of experiment with frequency and number of Foxp3^+^CD4^+^ T_reg_ cells in pre-onset tumor nodules. FVB mice received P5-control vector, MP5-control vector and P5-CCL2^OE^ MaSCs, and 20 or 10 days later pre-onset tumors were extracted and analyzed by flow cytometry. **(g)** Lethally irradiated CD45.1^+/-^ recipient mice were reconstituted with WT CD45.1^+/+^ and *Ccr2^KO^* CD45.2 BM. Six weeks post reconstitution, MP5-sg*NT* cells were injected to mice and 10 days later, pre-onset tumor nodules and dLNs were collected for FACS analysis. Graphs show the frequency and numbers of CD45^+^ immune, Foxp3^+^CD4^+^ T_reg_ and non-T_reg_ CD4^+^ T cells in WT and *Ccr2^KO^*hematopoietic-derived (CD45^+^) compartment of pre-onset nodules. Representative dot plots and summary plots from 2 experiments are shown (n=5). P-values were determined by unpaired t-test (a,b,c, and f) or Log-rank test (d and e) or paired t-test (g). *p< 0.05, **p< 0.01, ***p< 0.001, ****p< 0.0001, ns or not indicated: non-significant.

To further assess whether CCL2 contributes to tumor development, we treated mice implanted with MP5 MaSCs with anti-mouse CCL2 neutralizing or isotype control Abs (**Fig. 6d and Supp. Fig 5g**). While anti-CCL2 Ab treatment significantly delayed MP5 tumor onset by ∼10 days compared to isotype control treated group, tumor growth was comparable in both groups, indicating that CCL2 contributed to early tumor escape but not growth. Of note, however, the number of all infiltrating leukocytes (CD45^+^), including T_reg_ cells, CD4^+^ T_conv_ cells, CD8^+^ T cells, B cells and Ly6C^+^ monocytic cells in MP5 pre-onset tumor nodules of anti-CCL2 Ab-treated mice were lower compared to isotype-Ab treated mice, suggesting a broad impact of systemic CCL2 neutralization (**Supp. Fig. 5h**). Thus, to directly link CCL2 secreted by tumor cells to T_reg_ cell infiltration and faster tumor onset, we transduced P5 MaSCs with a *Ccl2*-expressing lentiviral vector to establish CCL2-overexpressing P5 MaSCs (P5-CCL2^OE^). CCL2 expression in P5-CCL2^OE^ tumors was comparable to CCL2 levels in MP5 tumors transduced with the empty lentiviral vector and ∼two-fold higher than control P5 tumors, as quantified by ELISA (**Supp. Fig. 5i**). P5-CCL2^OE^ MaSCs formed tumors significantly faster than control P5 cells but still at slower pace than control MP5 MaSCs (**Fig. 6e**). Consistent with our hypothesis, P5-CCL2^OE^ pre-onset tumors (day 10) had significantly higher number of infiltrating T_reg_ cells than P5 tumors (day 20), but similar numbers to MP5 tumors (**Fig. 6f**). Of note, while there was no significant difference in the overall number of tumor-infiltrating CD45^+^ cells (and NK, CD8^+^ T cells and Ly6G^+^ granulocytic cells), the number of CD4^+^ T_conv_ cells, Ly6C^+^ monocytic and F4/80^+^ MPs cells were significantly higher in P5-CCL2^OE^ pre-onset tumors (**Supp. Fig. 5j**).

Since CCL2 from tumor cells promoted higher numbers of tumor-infiltrating T_reg_ cells in pre-onset tumor nodules, we predicted that CD4^+^ T_reg_ cells lacking CCR2, the major receptor for CCL2, should fail to infiltrate the tumor nodules. To formally test this possibility, we generated mixed bone marrow (BM) chimeras in which WT B6 CD45.1^+/-^ mice were lethally irradiated and reconstituted with a 1:1 ratio of WT (CD45.1^+/+^) and *Ccr2*^-/-^ BM cells (**Fig. 6g and Supp. Fig. 5k**). The frequencies and numbers of WT and *Ccr2*^-/-^ hematopoietic-derived (CD45^+^) cells in blood and tumors were close to the reconstituting ratio (1:1). Most interestingly, the proportion and numbers of *Ccr2*^-/-^ T_reg_ cells were significantly lower than that of WT counterpart in tumor nodules while tumor-infiltrating *Ccr2*^-/-^ CD4^+^ T_conv_ cells were increased compared to WT counterparts. This result suggested that CCR2 significantly enhanced the competitive recruitment of T_reg_ cells to pre-onset MP5 tumor nodules. *Ccr2*^-/-^ T_reg_ cell proportions and numbers in dLNs were comparable to WT counterparts (**Supp. Fig. 5l**). Collectively, these data establish that CCL2 produced by tumor cells promote CCR2^+^ T_reg_ cell recruitment and infiltration to early stage pre-onset tumors, accelerating tumor escape. The partial rescue of P5-CCL2^OE^ tumor onset compared to MP5 tumors, further suggested that additional non-CCL2-dependent mechanisms are likely involved in accelerated MP5 tumor onset.

### HIF1α promotes the differentiation of effector T_reg_ cells with a highly immunosuppressive phenotype in early stage tumors

T_reg_ cells infiltrating MP5 pre-onset tumors acquire a highly immunosuppressive phenotype compared to P5 counterparts (**Fig. 3**). Thus, we hypothesized that in addition to regulating CCL2 secretion, HIF1α also alters the differentiation of CCR2^+^ infiltrating T_reg_ cells. We characterized T_reg_ cells infiltrating MP5-sg*Hif1a* and P5-CCL2^OE^ pre-onset tumors using our T_reg_ cell-focused spectral flow cytometry panel (**Table S1**). UMAP analysis of tumor-infiltrating T_reg_ cells revealed a comparable distribution of T_reg_ cells in P5 and P5-CCL2^OE^ pre-onset tumors but distinct from MP5 tumors. Interestingly, T_reg_ cells in MP5-sg*Hif1a* tumors segregated differently from both P5 and MP5 tumors, consistent with our hypothesis (**Fig. 7a**). Notably and compared to MP5 tumors, T_reg_ cells in MP5-sg*Hif1a* tumors expressed lower levels of markers associated with effector fate (KLRG1, Blimp-1), proliferative (Ki67), and suppressive (ICOS, GITR, CD25, CTLA-4) functions (**Fig. 7b and Supp. Fig. 6a**). Further characterization of T_reg_ cells in P5-CCL2^OE^ pre-onset tumors where the number of recruited T_reg_ cells is increased and comparable to that of MP5 tumors (**Fig. 6f**), showed that infiltrating T_reg_ cells mostly failed to differentiate into effector T_reg_ cells with a highly immunosuppressive phenotype, confirming that other HIF1α-dependent signals, are needed for this to occur in addition to CCL2/CCR2-mediated recruitment of T_reg_ cells. Interestingly, overexpression of CCL2 and lack of HIF1α impacted both tumor-infiltrating CD4^+^ T_conv_ and CD8^+^ T cell phenotypes, yet most changes except for loss of specific inhibitory receptors expression (ICOS, CTLA-4, PD-1), did not seem to be directly related to the loss of MLL3 (i.e., P5 versus MP5, **Supp. Fig. 6b, c**). Thus, altogether, these results suggest that HIF1α orchestrates both the recruitment (via CCL2/CCR2) and the differentiation of highly suppressive effector T_reg_ cells early on, allowing for rapid tumor cell escape and growth.

**Figure 7.**
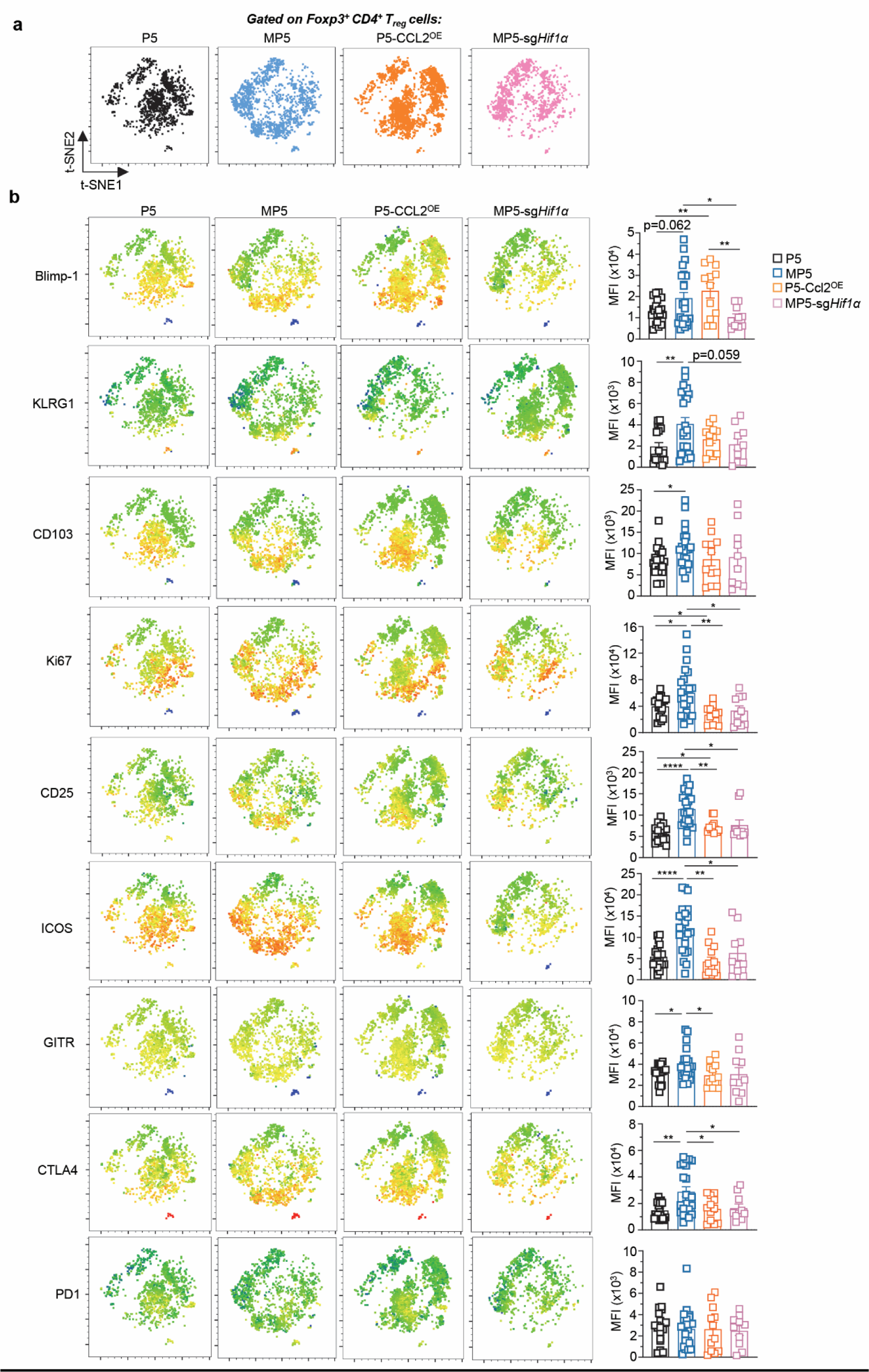
HIF1α promotes the differentiation of effector T_reg_ cells with a highly immunosuppressive phenotype in early stage tumors. **(a)** T-SNE spatial distribution of T_reg_ cells infiltrating indicated pre-onset tumors based on 22-marker flow cytometry analysis (Data of P5 and MP5 is same as in Figure 3). **(b)** Relative expression level of indicated markers at the single cell level on t-SNE maps with bar graphs averaging overall expression level for each marker in T_reg_ cells. t-SNE maps concatenate data from 4 independent replicate experiments, in P5 (n=19), in MP5 (n=22), in P5-CCL2^OE^ (n=13), in MP5-sg*Hif1α* (n= 9) tumors. Each symbol (bar graphs) feature 1 tumor. P-values were determined by unpaired t-test. *p< 0.05, **p< 0.01, ****p< 0.0001, ns or not indicated: non-significant.

### Therapeutic targeting of ICOS or GITR prevents tumor onset

Among the most upregulated cell-surface markers found on T_reg_ cells infiltrating MP5 compared to P5, P5-CCL2^OE^ and MP5-sg*Hif1a* tumors, we noted ICOS and GITR (**Fig. 6b**). While also expressed on both CD4^+^ and CD8^+^ T_conv_ cells in the tumors, their levels were significantly lower compared to that of T_reg_ cells (**Fig. 8a**). This led us to postulate that therapeutic targeting of these pathways may more selectively target T_reg_ cells to delay or prevent tumor onset. We therefore treated mice implanted with MP5 MaSC either before tumor onset or after developing established tumors (∼50mm^3^) with i) an ICOS antagonistic mAb (clone 7E17G9) that prevents ICOS/ICOSL interactions, ii) a GITR agonistic mAb (clone DTA-1) reported to deplete and/or functionally impair T_reg_ cells’ability to suppress^11,60–63^, prevented tumor onset while isotype-treated groups all had detectable tumors by day 30, consistent with our hypothesis (**Fig. 8b**). Moreover, eight weeks after each treatment ended, ∼80% (GITR) and 60% (ICOS) of the mice remained tumor free, suggesting a durable response. In addition, both treatments prevented the majority of established tumors to grow further (∼50-75%, **Fig. 8c**). GITR targeting was also more effective than ICOS targeting. Interestingly, in the TCGA BRCA dataset, we also observed a negative correlation between *KMT2C/MLL3* mRNA expression and that of both *ICOS* and *GITR* (**Fig. 8d**). Thus, therapeutic targeting of GITR, and possible ICOS, at early stage of tumor escape as well as on established tumors, may represent a promising therapeutic strategy for *Mll3*-mutant and HIF1α^hi^ breast cancers.

**Figure 8.**
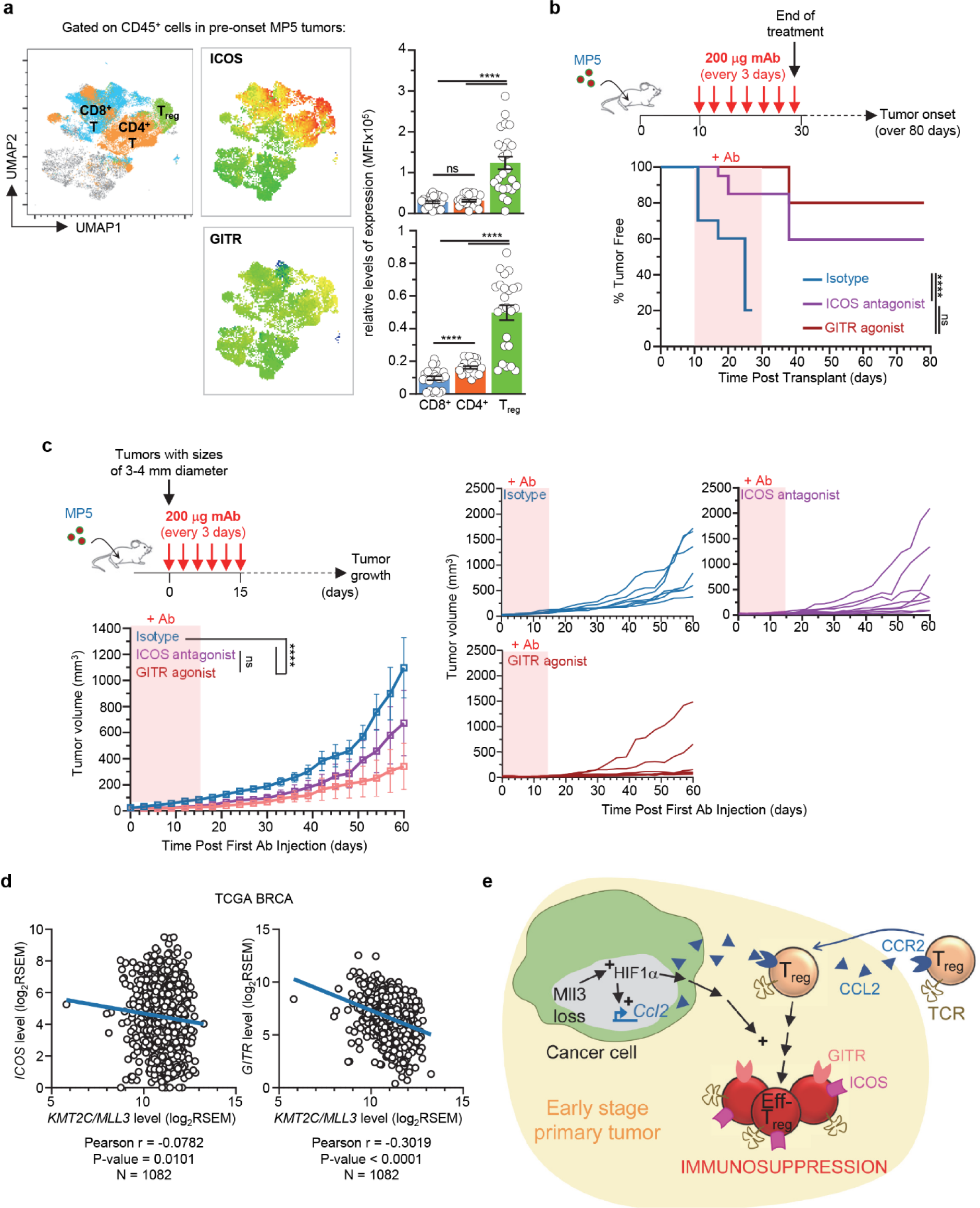
Therapeutic targeting of ICOS or GITR prevents tumor onset. **(a)** UMAP overlays of indicated T cell populations on tumor-infiltrating immune cells (CD45^+^) and relative cell-surface expression levels of ICOS and GITR distributed on the UMAP at the single-cell level. Bar graphs average overall CD8^+^ T, CD4^+^ T_conv_ and T_reg_ cells expression level for both markers. **(b)** Experimental design and Kaplan-Meier analysis of tumor onset in WT B6 mice implanted with MP5 MaSCs and treated with ICOS antagonist, GITR agonist or control isotype mAbs. Data are pooled from two independent replicate experiments (n=10 mice per group). **(c)** Experimental design and tumor growth in WT B6 mice implanted with MP5 MaSCs and treated with ICOS antagonist (n=8), GITR agonist (n=8) or control isotype mAbs (n=7). Right graphs show growth of individual tumors. **(d)** Correlation of KMT2C and ICOS or GITR gene expression in TCGA Pan Cancer Atlas Breast Cancer database (N=1082). RSEM, RNA-seq by Expectation Maximization. **(e)** Working model: loss of the new tumor suppressor gene *Mll3* activates HIF1α and CCL2 secretion from tumor cells, which promotes Foxp3^+^ T_reg_ cell i) infiltration via CCR2-CCL2 and ii) differentiation into effector ICOS^hi^GITR^hi^ effector T_reg_ cells (Eff-T_regs_). P-values were determined by unpaired t-test (a) or Log-rank test (b) or Two-way-ANOVA (c) or Pearson’s rank correlation (d). ****p< 0.0001, ns: non-significant.

## DISCUSSION

In this work, we show that the functional loss of the MLL3 tumor suppressor protein alters the immune response in the TME. MLL3 loss increased expression of the transcription factor HIF1α inside the tumor cells. HIF1α induction in tumor cells enhanced i) the infiltration of higher numbers of T_reg_ cells and ii) their differentiation into Blimp-1^+^ effector T_reg_ cells that co-express high cell surface levels of ICOS and GITR in the TME. Mechanistically, HIF1α appears to transcriptionally promote the secretion of the chemokine CCL2 by tumor cells through binding to the HRE-containing regulatory elements close to the *Ccl2* transcription start site. CCL2 from tumor cells then enhances the rapid infiltration of Foxp3^+^ T_reg_ cells via CCR2 in *Mll3*-mutant tumors. Consistent with a role during early-stage tumor escape, selective depletion of T_reg_ cells significantly delayed tumor onset (**Fig. 8e**). MLL3 mutation or downregulation in human breast tumor biopsy tissue sections also correlated with Foxp3^+^ T_reg_ cell infiltration. Very interestingly, therapeutic targeting of ICOS or GITR, prevented i) 60-80% of MLL3-mutant tumors from growing *in vivo* and ii) inhibited the growth of established tumors, offering a potential treatment opportunity for MLL3-mutant (and possibly also MLL3-WT) TP53/PIK3CA breast cancers.

MLL3 is frequently altered in human cancer, and emerging evidence has suggested its role in tumor suppression. However, how MLL3 and co-occurring cancer mutations alter the TME is not known. Using MaSC-based tumor models that combine the loss of MLL3 with PI3K activation and p53 knockout mutations, a genetic setting that occurs in human breast cancer, we show that functional loss of MLL3 accelerated tumor onset and growth indicating a potent tumor suppressor role of MLL3. We reveal that the loss of MLL3 lead to the upregulation of HIF1α, a key cancer signaling pathway. HIF1α expression correlates with poor clinical prognosis in many cancer types and contributes to a multitude of aggressive cancer behaviors, including proliferation, angiogenesis, metabolism, epithelial-mesenchymal transition, invasion, and cancer stemness^64,65^. HIFα (HIF1α and HIF2α) is often activated by the hypoxic tumor microenvironment in cancer, as 50 to 60% of solid tumors contain regions of hypoxia ^66^. Interestingly, recent studies also show that certain cancer cells exhibit high HIFα activity even under normoxia conditions^67,68^, suggesting that HIFα can also be regulated by mechanisms other than oxygen tension. The pre-onset tumor nodules are embedded in the well-oxygenated normal mammary tissue, therefore likely non-hypoxic. It is likely that, in the non-hypoxic pre-onset nodules, MLL3 loss upregulated HIF1α in a hypoxia-independent manner. Interestingly, in large mammary tumors which are often hypoxic, HIF1α is robustly expressed regardless whether MLL3 is expressed or not. Hypoxia is sufficient to induce robust HIF1α expression, bypassing the effect of MLL3 deletion. Agreeing with this result, MLL3 deletion increases T_reg_ cell infiltration in pre-onset nodules, but not in large tumors. The precise biochemical mechanism by which MLL3 regulates HIF1α still needs to be determined. Recent studies have shown that HIF1α can be methylated in several lysine residues by histone lysine methyltransferases SET7/9 and G9a, and these modifications affect the stability or transcriptional activity of HIF1α^69–72^. Future studies will be needed to determine whether MLL3 regulates HIF1α through similar mechanisms.

We established that MLL3-induced HIF1α enhanced CCL2 secretion by tumor cells, leading to CCR2-mediated recruitment of T_reg_ cells in the very early stages of tumor onset. Thus HIF1α, likely through direct regulation of CCL2, facilitates the establishment of an immunosuppressive microenvironment in early tumor lesions that are not yet hypoxic. These findings are consistent with an earlier report using the MMTV-PyMT model of breast cancer, where CCL2 from tumor cells was also reported to attract CCR2^+^ T_reg_ cells from the dLNs^18^. Similarly, CCR2 also contributed to T_reg_ cell recruitment to dLNs from inflamed tissues to suppress allograft rejection^73^. Therefore, CCR2 on T_reg_ cells from tumor dLNs are likely to enhance their access to Mll3-mutant tumors. While CD4^+^ T_conv_ cells may still migrate to the *Mll3*-mutant pre-onset tumor nodules via CCR2 and rapidly convert to T_reg_ cells, the proportion of CD4^+^ T_conv_ cells was comparable in both dLNs and pre-onset tumor nodules, and this ratio became inverted only for T_reg_ cells, rather suggesting direct recruitment of CCR2^+^ T_reg_ cells from the dLNs to the pre-onset tumor nodules. Consistent with this possibility, CCR2^-/-^ T_reg_ cells still infiltrated MP5 tumor nodules, albeit less efficiently than CCR2^+/+^ cells, suggesting the involvement of other chemotactic mechanisms. For instance, tumor hypoxia is reported to enhance T_reg_ cell recruitment through the induction of CCL28 expression^14^. T_reg_ cells are also shown to migrate in response to CCR4/CCL22 and accumulate in breast and ovarian carcinoma tumors, which is associated with worse clinical outcomes and relapse in patients^17,74^. Likewise, expression of Foxp3 transcripts correlates with that of CCR4/CCL22 in the TCGA tumor data set for colon, breast, pancreas and lung cancers ^75^. Transcriptomic analysis of tumor-infiltrating human T_reg_ cells in breast cancer compared to normal breast tissue-resident T_reg_ cells, shows an increased expression of genes encoding for multiple chemotactic receptors (CCR5, CCR8, CCR10, CXCR3, CX3CR1, CXCR6), which is also associated with more aggressive cancers^76^. Thus, while other chemotactic mechanisms distinct from CCL2/CCR2 likely contribute to the infiltration of T_reg_ cells in Mll3-mutant tumors, our data add to the abundant literature underlining the importance of the CCR2/CCL2 axis in cancer biology and its potential therapeutic promises^77^ ^78,79^.

Importantly, in addition to inducing CCL2, we discovered that HIF1α in *Mll3*-mutant tumor cells, enforces the differentiation of infiltrating T_reg_ cell into Blimp-1^+^KLRG1^+^ effector T_reg_ cells reported to exhibit highly immunosuppressive characteristics^80,81^ ^82,83^. T_reg_ cells infiltrating tumors in this model, express high cell-surface CD103 which is associated with tissue residency, and expand (Ki67^+^), possibly in response to tumor antigens. The fact that they upregulate high-level cell surface ICOS and GITR, together with other key checkpoint inhibitory molecules (PD-1, TIGIT, LAG-3), is also consistent with a very potent suppressive capacity^11,84^. Loss of MLL3 further enhanced tumor T_reg_ cells acquisition of activated (CD44^hi^CD62L^low^Ki67^hi^), suppressive (ICOS^hi^GITR^hi^), effector (KLRG1^+^) and residency (CD103^hi^) features. This observation further suggests that depending on the combination of cancer driver mutations, intratumoral T_reg_ cells will acquire different phenotypes. Overexpression of CCL2 in P5 tumors failed to promote the emergence of such effector T_reg_ cells, ruling out a direct role for CCL2 signaling in this process of T_reg_ cell differentiation. How exactly HIF1α expression by tumor cells orchestrates the differentiation of effector T_reg_ cells is unclear. HIF1α upregulation increases glycolysis and lactic acid production in the TME^85,86^. Thus, T_reg_ cells in HIF1α-high tumors may uptake and metabolize lactic acid to maintain proliferation and suppressive functions^21^.

Tumor infiltrating T_reg_ cells contribute to immunosuppression through a variety of mechanisms. We reveal that blocking ICOS/ICOSL interaction or triggering GITR signaling respectively prevented ∼60% and 80% of Mll3-mutant tumor onset, even ∼3 months after we arrested the therapeutic mAb treatment. ICOS, a key member of the CD28 co-stimulatory molecules, is very highly expressed on the Mll3-mutant tumor infiltrating T_reg_ cells. Blocking ICOS signaling is known to inhibit ICOS^+^ T_reg_ cell suppressive activity, possibly through preventing them to secrete immunosuppressive IL-10^60,87^. ICOS is also induced on Ag-activated CD4^+^ and CD8^+^ T cells, and its triggering is required to enhance antitumor T cell responses. It is therefore likely that our antagonistic mAb treatment targets tumor infiltrating T_reg_ cells rather than T_conv_ cells. Inhibiting the ICOS pathway was also reported to synergize with anti-CTLA-4 mAb treatment ^88^, which could further potentiate treatment efficacy in our model. With regards to GITR agonistic mAb treatment, a member of the TNF receptor family, it is well established that its stimulation abrogates T_reg_ cell-mediated immunosuppression both *in vitro* and *in vivo* while enhancing antitumor CD4^+^ T_conv_ and CD8^+^ T cell-responses ^61,62,89^. The remarkable impact of GITR stimulation treatment regimen on preventing *Mll3*-mutant tumor onset seems consistent with a direct effect on T_reg_ cells that also synergizes with a tumor-specific effector T cell responses. In summary, these results support our model in which early *Mll3*-mutant tumor infiltrating T_reg_ cells contribute to the rapid establishment of an immunosuppressive TME via ICOS and GITR, allowing faster tumor cell escape and growth.

In summary, our work demonstrates a major role of MLL3 loss, in combination with the most frequently associated breast cancer driver mutations *TP53* and *PIK3CA*, in promoting rapid mammary tumor onset and growth through the establishment of an immunosuppressive TME. This occurs via enhanced infiltration of Foxp3^+^ T_reg_ cells, a finding we also observed in human breast cancer biopsies. The fact that the HIF pathway is a key checkpoint downstream of the loss of MLL3, suggest a potentially generalizable mechanism to other, non-*MLL3*-mutant hypoxic tumors. CCL2 is also an essential chemokine for the recruitment of multiple types of pro-tumor immune cells^77^. Together with the effectiveness of ICOS and GITR targeting, our findings offer potentially novel therapeutic strategies for *MLL3*-mutant and hypoxic tumors.

## EXPERIMENTAL PROCEDURES

### Ethics statement

This study was carried out in strict accordance with the approvals by the animal use committee at the Albert Einstein College of Medicine. All efforts were made to minimize suffering and provide humane treatment to the animals included in the study. Human sample studies (IRB# 2018-9432) were approved by the Institutional Review Board of Albert Einstein College of Medicine.

### Mice

All mice were bred in the SPF animal facility at the Albert Einstein College of Medicine. Wild-type (WT) C57BL/6J (B6) (JAX, #000664), FVB/n (JAX, #001800), Pik3ca* (JAX, #012343), Pik3ca^H1047R^ (JAX, #016977), *Rag1*^-/-^ (JAX, #002216), *Foxp3*^DTR/GFP^ (JAX, #016958) and *Ccr2*^KO^ (JAX, #004999) mice were purchased from the Jackson Laboratory.

### Public database analyses

For analysis of cancer driver mutation frequency (Figure 1a), MSKCC-IMPACT 2017 Breast Cancer database was used in the cBioportal website. For determination of MP5-like breast tumor subtype (**Supp. Fig. 1a**), the TCGA BRCA cohort were analyzed in the cBioportal website.

For analyses of overall survival of P5 and MP5-like patients, *HIF1A* target genes and *CCL2* mRNA expression between MLL3 WT and mutant breast cancer patients, and Gene Set Enrichment Analysis (GSEA) of transcriptome from P5 and MP5-like patients, the dataset from the TCGA BRCA cohort (PanCancer Atlas, 1084 patients) was subject to analysis process via cBioPortal and GSEA (4.2.3).

### TIMER2.0 analyses and human breast cancer patient immunostaining

The correlation between MLL3 mRNA expression and T_reg_ cell infiltration in breast cancer patients (n=1100) was determined by TIMER2.0 with CIBERSORT algorism^90^.

Human breast cancer patient FFPE-tissue microarray (TMA) was purchased through TissueArray.com LLC (#BC081116e). The slide with sample cores was stained with antibodies against Pan-Keratin, FOXP3 and MLL3. To reduce background signal in MLL3 staining, which comes from non-specific binding, the MLL3 antibody was pre-absorbed by MP5 MaSCs. Briefly, MP5 MaSCs (5×10^5^) were seeded in 24-well normal cell culture plate without Matrigel to grow in an adhesion condition. Two days after the cells were seeded (95% confluent), the cells were washed with PBS, and fixed with 4% PFA. The fixed cells were subjected to 0.2% Tritons X-100 in PBS for permeabilization, and incubated with blocking solution (2% BSA+5% normal goat serum+0.1% Tween 20 in PBS). After blocking, the cells were incubated with MLL3 antibody diluted in staining buffer (5% BSA+5% normal goat serum+0.1% Tween 20 in PBS) for overnight. The pre-absorbed antibody was then used for TMA staining. The stained slides were scanned by 3DHistec Pannoramic 250 Flash II slide scanner, and the number of FOXP3^+^ cells and the mean nuclear MLL3 intensity in tumor cells (Keratin^+^) was counted and measured by CellProfiler4.2.5 ^91^. The number of FOXP3^+^ cells were normalized by non-tumor area.

Fifty-seven fresh-frozen breast cancer tumor (T) samples and matching non-tumor (NT) tissues were obtained through IRB# 2018-9432, and were analyzed for mutational profiling using the Einstein Custom Cancer Panel (ECCP), a mid-size cancer focused targeted next generation sequencing panel that includes 156 oncogene and tumor suppressor genes commonly mutated in human tumors ^92^, including *TRP53*, *PI3KCA* and *KMT2C* genes. From the cohort we identified 4 samples carrying pathogenic mutations both in TP53 and PIK3CA, two with wild type KMT2C (#542 and #586) and two with KMT2C mutations (#125 and #354) (see Table S2). (see Table S2). The frozen sections were fixed with cold acetone and incubated with 0.05% TBS-Tween 20 (TBS-T), followed by incubation with 1% TritonX-100 in TBS. After blocking with 5% BSA and 5% goat serum in TBS, the slides were stained with anti-FOXP3 antibody (see Table S1). After primary antibody overnight incubation, the slides were stained with secondary antibodies and counterstained with DAPI. The stained slides were scanned by 3DHistec Pannoramic 250 Flash II slide scanner, and the number of FOXP3^+^ cells was counted by CellProfiler4.2.5 ^91^. The number of FOXP3^+^ cells were normalized by DAPI^+^ area measured by CellProfiler.

### Isolating MaSCs and culturing MaSC organoid

Mammary glands were minced and digested with 300 units/ml collagenase type 3 (Worthington Biochemical, LS004182), and 10 μg/ml Dnase I (Worthington Biochemical, LS002139) in the DMEM/F-12 medium at 37°C for 2 hrs on an orbital shaker (100 units/ml hyaluronidase added only for tumor digestion). The digested samples were washed with PBS and were further digested with 0.05% trypsin-EDTA for 5 minutes and 1 unit/ml neutral protease (dispase) (Worthington Biochemical, LS02109) plus 100 ug/ml Dnase (Roche) for 5 minutes. The digested cells were then filtered through 40 μm cell strainers to obtain single cells.

Organoid culture was performed as previously reported ^93^. Organoid medium was prepared with Advanced DMEM/F-12 (Life Technologies) supplemented with 5% heat-inactivated FBS (Sigma, F2442), 10 ng/ml EGF (Sigma, E9644), 20 ng/ml FGF2 (EMD Millipore, GF003), 4 μg/ml heparin (Sigma, H3149), 5 μM Y-27632 (Cayman Chemical, #10005583) and 5% Matrigel (Corning, #354234). For passaging and expansion, organoids were washed with PBS, dissociated with 0.05% trypsin/EDTA and seeded at 3 × 10^5^ cells/well in 6-well ultra-low attachment plates. Cells were then passaged every 3-4 days. Dissociated organoid cells were frozen in calf serum containing 10% DMSO and 5 μM Y-27632, and thawed to reestablish the culture.

### Viral transduction

Lentiviruses were produced with the pMD2.G and pCMVR8.74 packaging system, a gift from Didier Trono (Addgene, plasmids #12259 and #22036). Viruses were concentrated with the Lenti-X Concentrator reagent (Clontech, #631232). For viral transduction, dissociated organoid cells were seeded in virus-containing organoid culture media with 5 μg/ml polybrene (EMD-Millipore TR-1003-G) overnight, and medium was refreshed the next day. Transduced cells were either FACS sorted by GFP or RFP (70-90% transduction efficiency) or selected by puromycin (2 μg/ml) or blasticidin (10 μg/ml) treatment.

### CRISPR-based genome editing

Genome editing for Pik3ca^H1047R^ MaSCs (FVB/n background) were carried out as following: MaSCs (passage 2-3) were transduced with sgP53 or sgNT cloned into lentiCRISPRv2-puro (Addgene, Plasmid #98290), passaged once, and then transduced with sgMll3 or sgNT cloned into pLKO5.sgRNA.EFS.tRFP (Addgene, plasmid #57823). Cells were grown for 3 days and selected with puromycin for 3 days. Next, cells were infected with Adeno-Cre-GFP and sorted for GFP/RFP double positive cells. MP5-Hif1α KO lines were made via transducing MP5 cells with sgHif1a or sgNT cloned into lentiCRISPRv2-GFP (Addgene, plasmid #82416) and sorting for GFP.

Genome editing for Pik3ca* MaSCs (B6 background) were carried out as following: MaSCs (passage 2) were transduced with sgP53 or sgNT cloned into lentiCRISPRv2-puro, passaged once and infected with Ad-Cre-GFP, and puromycin selected for 3 days. Cells were then transduced with sgMll3 or sgNT cloned into pLKO5.sgRNA.EFS.tRFP and sorted for GFP/RFP double positive cells.

Targeting sequences for sgRNAs are as following: sgMll3 GCACACGATCTAGTACTCAG (Exon10), sgp53 CCTCGAGCTCCCTCTGAGCC (Exon 2), sgNT GCGAGGTATTCGGCTCCGCG, and sgHif1a CACAATGTGAGCTCACATCT (Exon 2).

### Overexpression of CCL2 in MaSC

The full-length cDNA of mouse *CCL2* (NM_011333.3) was amplified by PCR with PrimeSTAR Max DNA Polymerase (Takara Bio, #R045A) from mouse MCP-1/CCL2 cDNA ORF Clone (Sino Biological, # MG50368-UT). The amplified insert was subcloned into pLV-EF1α-IRES-Blast vector (Addgene, plasmid #85133), which is then used to for generating P5-CCL2^OE^. For the controls, the original empty vector was used for virus production (P5-empty and MP5-empty). The line#1 P5 and MP5 MaSCs were transduced and selected by blasticidin (10 μg/ml). The CCL2 levels in tumor lysate was measured by MCP-1 Mouse Uncoated ELISA Kit (Thermo Scientific, #88-7391-22) with comparable sized tumors (∼3 mm diameter), according to the manufacturer’s protocol.

### Tumor onset and growth

Organoids were cultured for 3-4 days, then dissociated and passed through 40μm cell strainer to obtain single cells. Single cells (3 x 10^5^) suspended in PBS and 25% Matrigel were orthotopically injected per #3 or #4 mammary gland of B6 or FVB WT mice aged 6-8 weeks. For tumor onset and growth experiments, 2 glands per mouse were injected. Specifically, P5 and MP5 lines #1, #3, #4 and Hif1a-KO and CCL2-OE lines were transplanted to FVB mice and lines #2 and #7 were transplanted to B6 mice. Tumor onset was monitored by weekly palpation, and Kaplan-Meier survival analysis was used to compare tumor onset. Tumor onset was scored when at least one dimension of the tumor measured ≥3mm. Tumor volume was measured by caliper and calculated as V = 0.532 × Length × Width^2^. All experiments done on pre-onset nodules (≤ 3mm or 20mm^3^) were extracted on 20 days for P5 and 10 days for MP5 post transplantation due to the ∼2-fold difference in growth rate between these two tumor types.

### Immunostaining on tissue sections and western blot

Formalin-fixed paraffin-embedded (FFPE) sections for organoids or tumors were stained with antibodies against HIF1α, Ki67, cleaved-caspase 3, Foxp3, H3K4me1, H3K4me2, and H3K27ac. Western blots were performed with antibodies against MLL3, phospho-Akt (Ser473), HIF1α, H3K4me1, H3K4me2, H3K27ac, Histone H3, β-actin, and α-tubulin (see Table S1 for antibody sources and dilutions).

### RT-qPCR

Total RNA was isolated from P5 and MP5 MaSCs by *Quick*-RNA Microprep Kit (Zymo Research, #R1050), and subjected to cDNA synthesis by High-Capacity cDNA Reverse Transcription Kit (ThermoFisher, #4368814), according to the manufacturer’s protocol. Quantitative PCR was performed with PowerUp SYBR Green Master Mix (ThermoFisher, #A25742), according to the protocol provided with the kit. See Table S3 for primer sequences.

### ChIP-qPCR

After one day of 5% O_2_ hypoxia culture, MP5 line#3 cells were crosslinked with 1% formaldehyde at room temperature based on ENCODE protocol ^94^. Crosslinked cells were subjected to the sonication process with Bioruptor (30 sec ON/30 sec OFF, 60 cycles, High speed), and the fragmentation was validated by electrophoresis in a 1% agarose gel. 30 µg chromatin was used for each immunoprecipitation (IP) and 5 µg chromatin was used for input. Briefly, 200 µL of Dynabeads M-280 Sheep Anti-Mouse IgG (Invitrogen, #11201D) washed with wash buffer (5 mg/mL BSA in PBS) was mixed with 1 µg Hif1a Ab (Novus Biologicals, NB-100-105) for 4 hrs at 4℃ rotating. After washing antibody-bound beads in 100 mM Tris-HCl pH 7.5, 500 mM LiCl, 1% NP-40, 1% sodium deoxycholate, the beads were resuspended with fractionated chromatin by magnetic stand, mixed for 1 hr at room temperature, then 3 hrs at 4℃ rotating. The chromatin conjugated beads were further washed with LiCl IP wash buffer 4 times, following wash with TE buffer (10 mM Tris-HCl pH 7.5, 0.1 M Na2EDTA). The washed chromatin conjugated beads were incubated at 65℃ overnight to reverse crosslink. ChIP DNA and Input DNA were purified with ChIP DNA Clean & Concentrator (Zymo Research, #D5205). Quantitative PCR was performed with PowerUp SYBR Green Master Mix, according to the protocol provided with the kit. See Table S3 for primer sequences.

### Preparation of cell suspensions for flow cytometry and adoptive transfers

Tumors were minced into 2-4 mm fragments, which were incubated in HBSS medium with 10 UI/mL collagenase I, 400 UI/mL collagenase D and 30 µg/mL at 37℃ for 1 hr under agitation. After, the fragments were homogenized, filtered through a 100µm nylon mesh cell strainer and cells were centrifuged at 1,450 rpm for 10 min at 4℃. Draining lymph nodes were incubated in HBSS medium with 400 UI/mL collagenase D at 37℃ for 1 hr under agitation. BM cells were obtained by flushing femur with complete medium (RPMI 1640, 10% FBS, 1% Penicillin/Streptomycin, 55 μM β-mercaptoethanol, 1 mM Sodium Pyruvate, 1X Glutamax, 1X non-essential amino acids) containing 10% FCS.

### Cell-staining for FACS analysis

Cell suspensions were incubated with 2.4G2 Fc Block Ab and stained in PBS 1%FCS, 2mM EDTA, 0.02% sodium azide with fluorescently tagged Abs purchased from eBioscience, BD Biosciences, R&D systems, or BioLegend (See Tables S1). Brilliant stain buffer (BD) was used when more than two Abs were conjugated with BD Horizon Brilliant fluorescent polymer dyes. Cells were stained for cell-surface marker expression, then fixed in eBioscience Fixation/Permeabilization buffer prior to intracellular granzyme B and Foxp3 transcription factor (TF) staining in eBioscience Permeabilization buffer for 1hr. Data acquisition was done using a FACSAria III or a Cytek Aurora flow cytometer. All flow cytometry data were analyzed using FlowJo v9 or v10 software (TreeStar).

### Multiplex-beads cytokine array assay

A custom-made mouse Cytokine 26-Plex-assay kit was designed and purchased from Invitrogen. The kit included specific quantification of mouse Eotaxin, IL-1α, IL-1β, IL-2, IL-3, IL-4, IL-5, IL6, IL-7, IL-9, IL-10, IL-12p70, IL-13, IL-15/IL-15R, IL25, CCL2, CCL3, CCL4, CCL5, CXCL2, CXCL10, TNFα, IFN-γ, GM-CSF, VEGF, GM-CSF, MCSF. Lyophilized standards were reconstituted with kit Assay Diluent and combined to 1 mL for stock solution and serially diluted to generate standard curves for each cytokine. Samples were diluted with Assay Diluent and the assay was performed in a 96-well filter plate using the kit’s components, according to the manufacturer’s protocol.

### *In vivo* T_reg_ cell depletion and mAb treatments

#### Treg cell depletion

Foxp3^DTR^/^GFP^ male B6 mice were crossed with FVB female mice, and the B6/FVB F1 mice were crossed with Foxp3^DTR^/^GFP^ male B6 mice again to obtain Foxp3^DTR^/^GFP^ homozygous female F2 mice (75% B6 background). The F2 mice were treated Diphtheria Toxin (Calbiochem, # 322326) (6 ng per gram of mouse body weight) twice every week on consecutive days.

#### mAb treatments

Anti-mouse MCP-1/CCL2 (clone 2H5) and hamster IgG2a isotype-control were purchased from BioXCell (USA). 3 days post MaSCs injection, treatment with anti-CCL2 or isotype Ab at 1mg/kg was initiated and repeated every 3 days across the course of the experiment. Anti-mouse ICOS (clone 7E.17G9), anti-GITR (clone DTA-1) and rat IgG2b isotype control were purchased from BioXCell (USA). For the tumor onset experiment, 10 days post MaSCs injection, treatment with anti-ICOS, anti-GITR or isotype Ab (200µg/mouse, IP) was initiated and repeated every 3 days for 20 days. Tumor onset was monitored for 2 months post treatment arrest. For the established tumor experiment, after the tumors reached size of 3-4 mm diameter, the treatment with anti-ICOS, anti-GITR or isotype Ab (200µg/mouse, IP) was initiated and repeated every 3 days for 15 days.

### Generation of bone-marrow chimera mice

Lethally irradiated (1,200 rads) WT B6 female mice were immediately reconstituted with a total of 2×10^6^ BM cells isolated from C*cr2^KO^* and WT at a 1:1 ratio. Chimerism of reconstituted mice was checked ∼6-8 weeks later in the blood, prior to MaSCs injection.

### Statistical analysis

P values were calculated using unpaired t-test except unless otherwise specified in figure legends. Log-rank test or Two-way-ANOVA was used in tumor onset and growth analysis. Spearman’s rank correlation test was used in TIMER2.0 analysis. The FWER-p value in GSEA was calculated with “classic” gene ranking and “Diff_of_class” setting. Paired t-test was used in bone marrow chimeric assay. Pearson’s rank correlation was applied in correlation analyses (*HIF1A* expression vs *CCL2* expression, and MLL3 intensity vs number of T_reg_). Welch’s t-test was used in median tumor onset comparison. All statistical analyses except for TIMER2.0 and GSEA were done in GraphPad Prism 9. P < 0.05 was considered significant (*p< 0.05, **p< 0.01, ***p< 0.001, ****p< 0.0001, ns or not indicated: non-significant).

## Acknowledgments

We thank the Einstein FACS, Histopathology, Analytical Imaging, and Stem Cell Isolation core facilities.

## Author contribution

MB, KN and PE designed, performed and interpreted a majority of experiments, assembled all Figures, and contributed to writing/editing of the manuscript. MS conducted HIF1α-binding site analysis and advised the ChIP experiments. EG conducted and analyzed the Luminex assays. EB contributed to the early characterization of immune cell tumor infiltrates. NC set up the high-dimensional FACS panels. ZZ developed and analyzed the initial P5/MP5 tumor models. CM established human breast cancer biopsies with cancer driver mutation data and advised on patient tissue analyses. MCdM contributed to patient tissue immunostaining. GL and WG designed and interpreted experiments with all authors, contributed to Figure design and editing, and wrote the paper.

## Data materials and availability

All data is available in the main text or the supplementary materials.

## Supplementary Figure and Table Legends

**Supplemental Figure 1, related to Figure 1.**
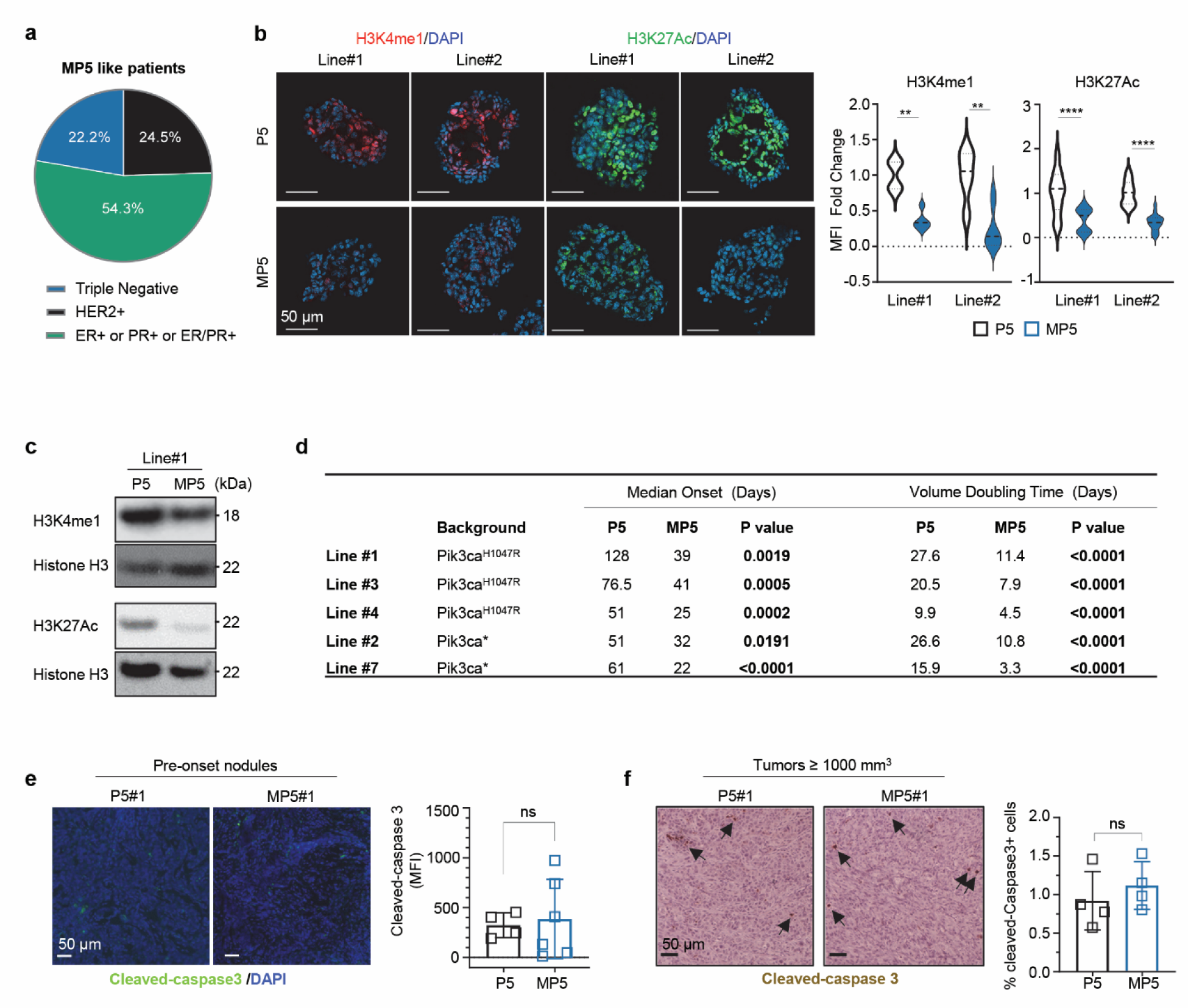
**(a)** Percentage of breast cancer with simultaneous *PIK3CA/MLL3/TP53* mutations in each breast cancer subtype (n= 63; TCGA, MSKCC-IMPACT and METABRIC databases). **(b)** Immunofluorescence images and quantification of H3K4me1 (top) and H3K27ac (bottom) levels in P5 and MP5 organoids from lines #1 and #2 (n= 15-30 organoids). **(c)** Western blot analysis for H3K4me1 and H3K27ac levels in P5 and MP5 organoids from line #1. **(d)** Medium tumor onset time and tumor growth rate for five pairs of P5 and MP5 MaSC models (n= 8-12 tumors per genotype of each line). 300K single cells were injected per #4 gland. **(e)** Immunofluorescence images and quantification of cleaved-caspase 3 expression in P5 (n=4) and MP5 (n=6) pre-onset tumor nodules (line #3). **(f)** Immunohistochemistry images and quantification of cleaved-caspase 3 expression in P5 (n=4) compared to MP5 (n=4) tumors (> 1000 mm^3^, line #1. P-values were determined by unpaired t-test (b,e, and f) or Log-rank test (d). **p< 0.01, ****p< 0.0001, ns: non-significant.

**Supplemental Figure 2, related to Figure 2.**
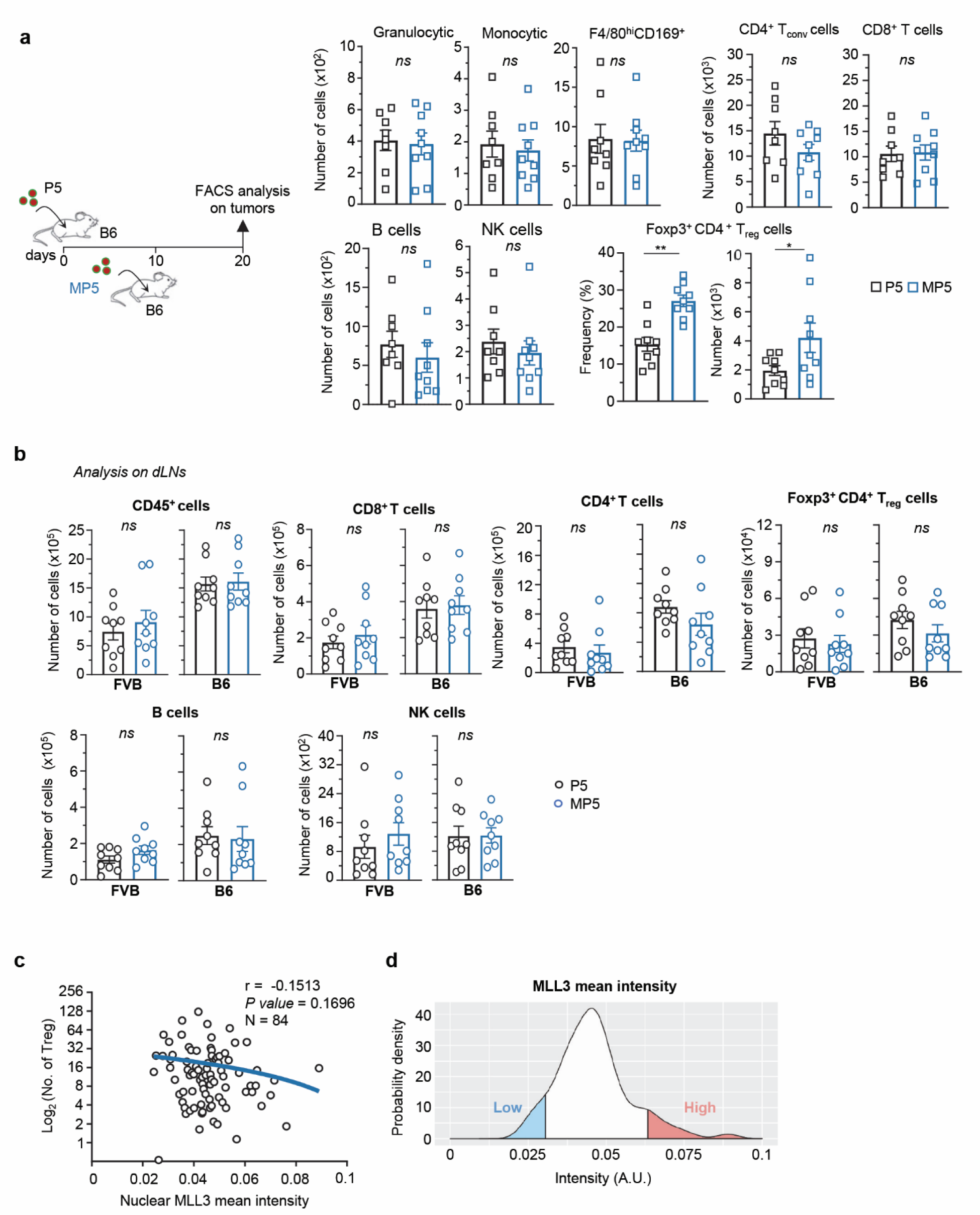
**(a)** Experimental design (WT C57BL/6 background). Numbers of indicated tumor infiltrating immune cells are shown. **(b)** Based on Fig. 2a and Suppl. 2b experimental designs: Numbers of CD45^+^ immune cells and indicated subsets in tumor draining lymph nodes of mice with P5 and MP5 pre-onset tumors. Data are from 2 independent replicate experiments (n=9 pre-onset tumor nodules). **(c)** Correlation between nuclear MLL3 mean intensity in pan-cytokeratin^+^ tumor cells and number of FOXP3^+^ T_reg_ cells in breast cancer patients by immunofluorescence analysis in TMA (n=84 cases). **(d)** Probability density curve for MLL3 intensity. Top and bottom 10% samples are highlighted in color. P-values were determined by unpaired t-test (a and b) or simple linear regression (c). *p< 0.05, **p< 0.01, ns: non-significant.

**Supplemental Figure 3, related to Figure 3.**
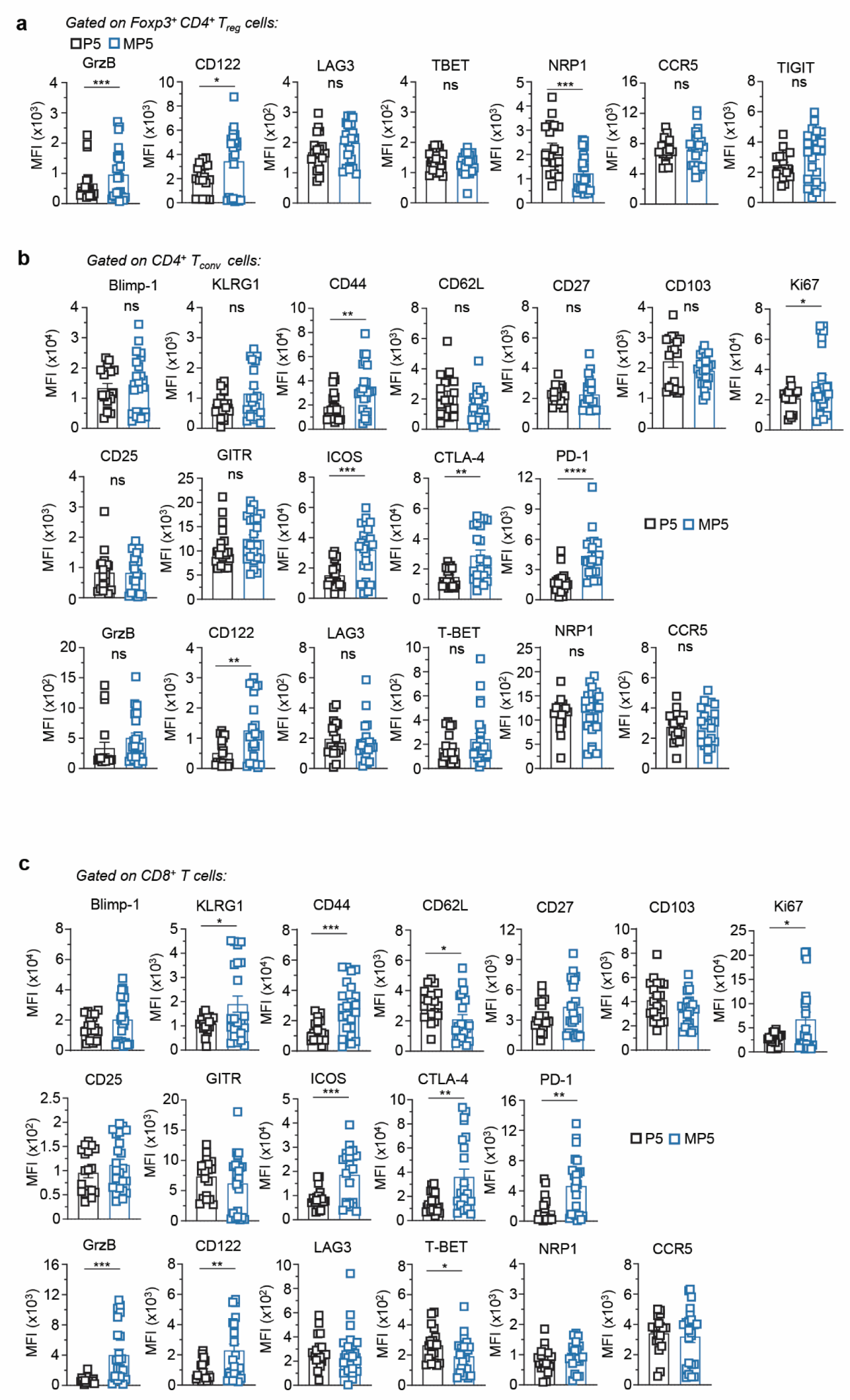
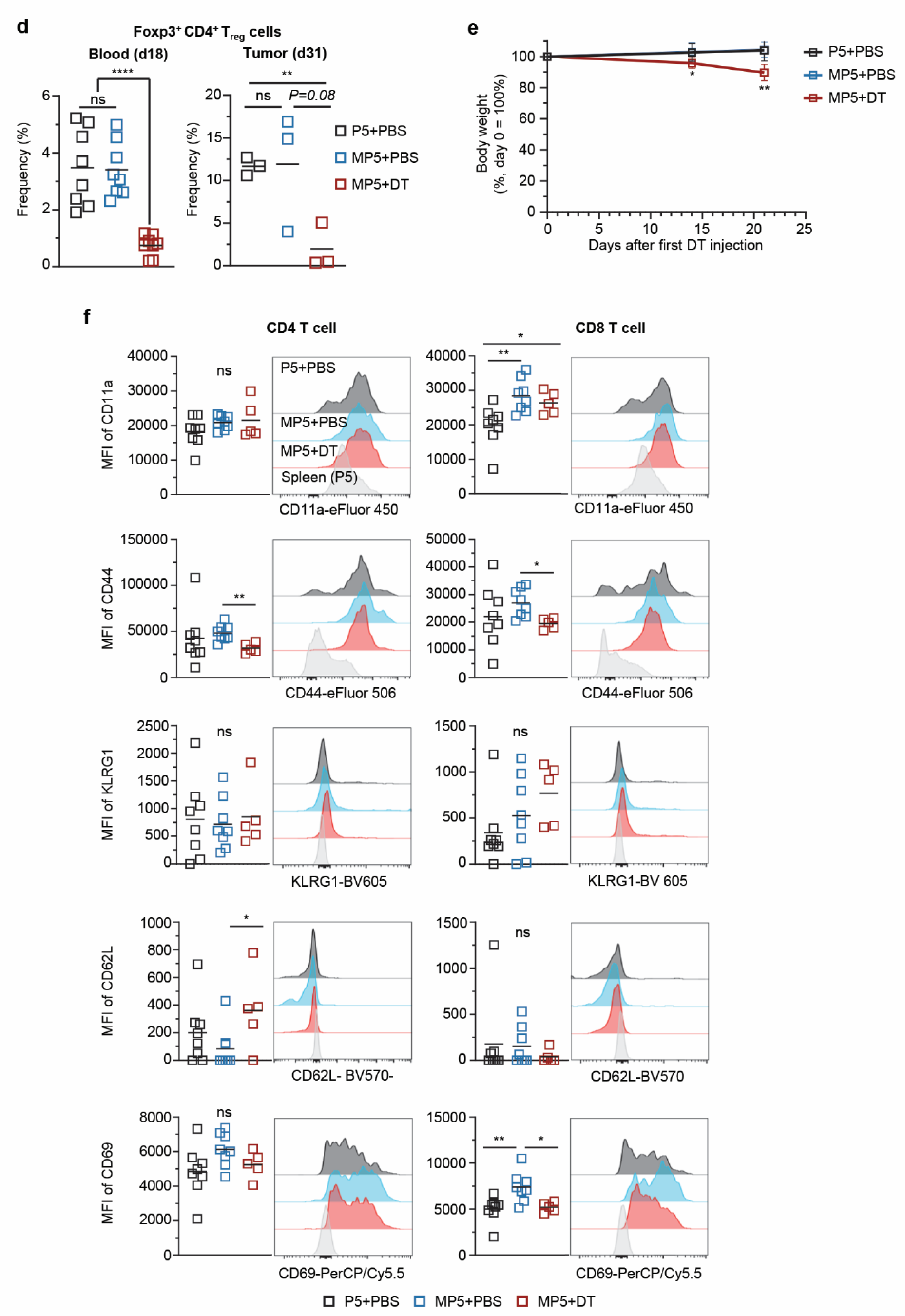
Bar graphs average the overall expression level for indicated markers on T_reg_ cells **(a)**, CD4^+^ T_conv_ cells **(b)** and CD8^+^ T cells **(c)** after staining of tumor infiltrating leukocytes with a 22 color marker panel. **(d)** Frequency of Foxp3^+^ CD4^+^ T_reg_ cells in the blood (day 18) and the endpoint tumors (day 31) (n=8 per group for blood, and n=3 per group for tumors). **(e)** Relative body weights, which are normalized to that at day 0 after first DT injection. **(f)** Expression levels (MFI) of indicated T cell activation markers in tumor-infiltrating CD4^+^ T_conv_ and CD8^+^ T cells in tumors (P5+PBS=8, MP5+PBS=8, MP5+DT=5). P-values were determined by unpaired t-test (a-f). *p< 0.05, **p< 0.01, ***p< 0.001, ****p< 0.0001, ns or not indicated: non-significant.

**Supplemental Figure 4, related to Figure 4.**
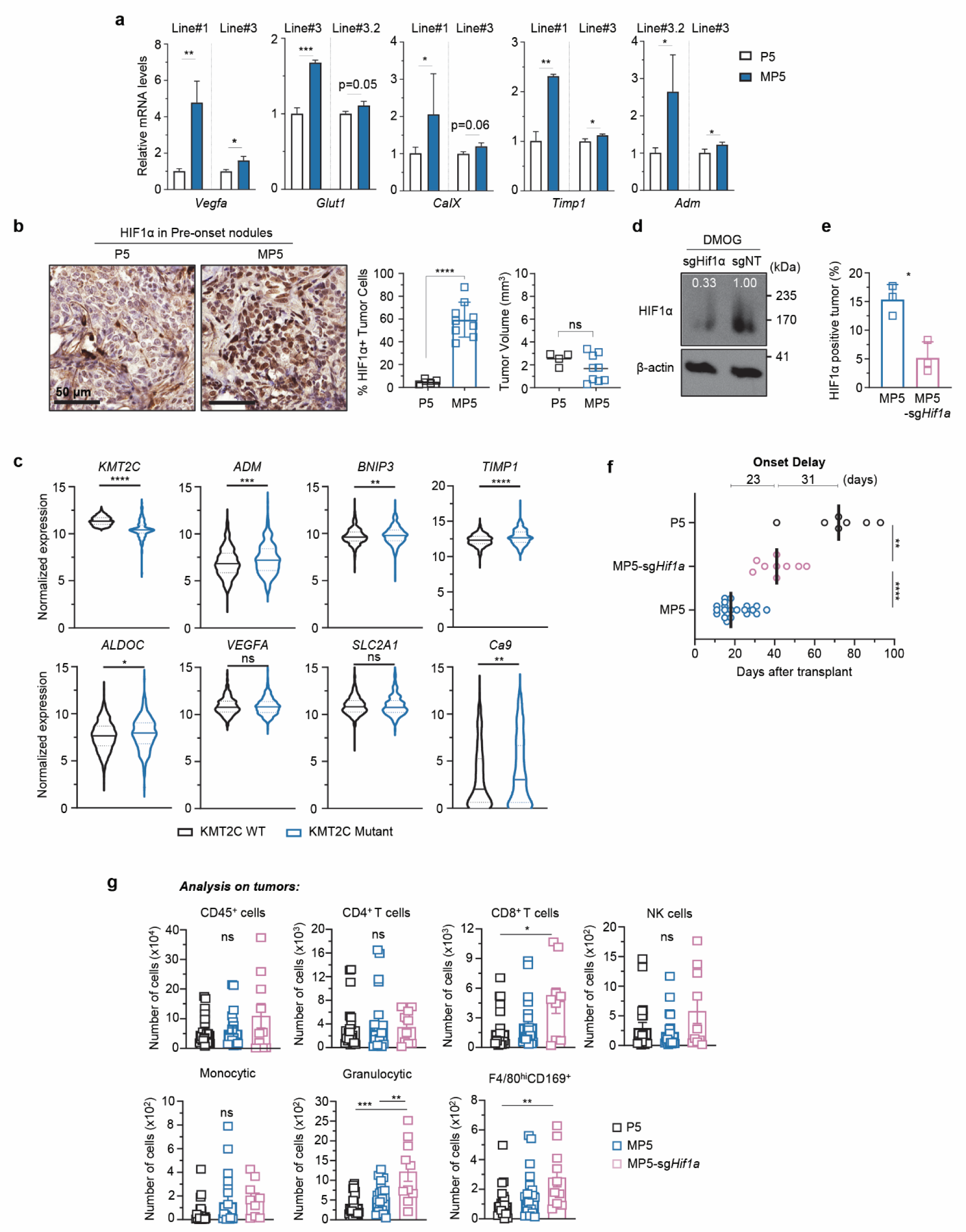
**(a)** mRNA expression of HIF1α downstream targets in P5 and MP5 organoids from lines #1, #3 #3.1 or #3.2. **(b)** Immunohistochemistry (IHC) images and quantification of HIF1α expression in P5 pre-onset tumor nodules (n=4) compared to MP5 (n=9) pre-onset tumor nodules from line #3. Volumes of tumor nodules used for HIF1α expression analysis were shown on the right. **(c)** *KMT2C*, *HIF1A* and *HIF1A* downstream targets mRNA expression levels in *MLL3*-WT (n=769) and *MLL3*-mutant (n=315) tumors in the TCGA Pan Cancer Atlas Breast Cancer database. **(d)** Western blot analysis of HIF1α expression in MP5-sg*NT* and MP5-sg*Hif1α* organoids treated with 1 mM DMOG for six hours. The values show relative band intensity of HIF1α normalized to that of β -Actin. **(e)** Quantification of HIF1α immunofluorescence staining images in RFP^+^ MP5-sg*NT* (n=3) or MP5-sg*Hif1α* (n=3) pre-onset tumor nodules. The values show the percentage of HIF1α^+^ tumors cells (RFP^+^) in total tumor cells. **(f)** Tumor onset time of P5-sg*NT* tumors (n=7), MP5-sg*NT* tumors (n=21) and MP5-sg*Hif1α* tumors (n=9), pool of two independent experiments. **(g)** Numbers of CD45^+^ and indicated cell subsets infiltrating P5-sg*NT*, MP5-sg*NT* and MP5-sg*Hif1α* pre-onset tumor nodules. Data are pooled from 3 independent experiment (n=9 samples/group). P-values were determined by unpaired t-test (a,b,c,e, and g) or Welch’s t-test (f). *p< 0.05, **p< 0.01, ***p< 0.001, ****p<0.0001, ns: non-significant.

**Supplemental Figure 5, related to Figure 5.**
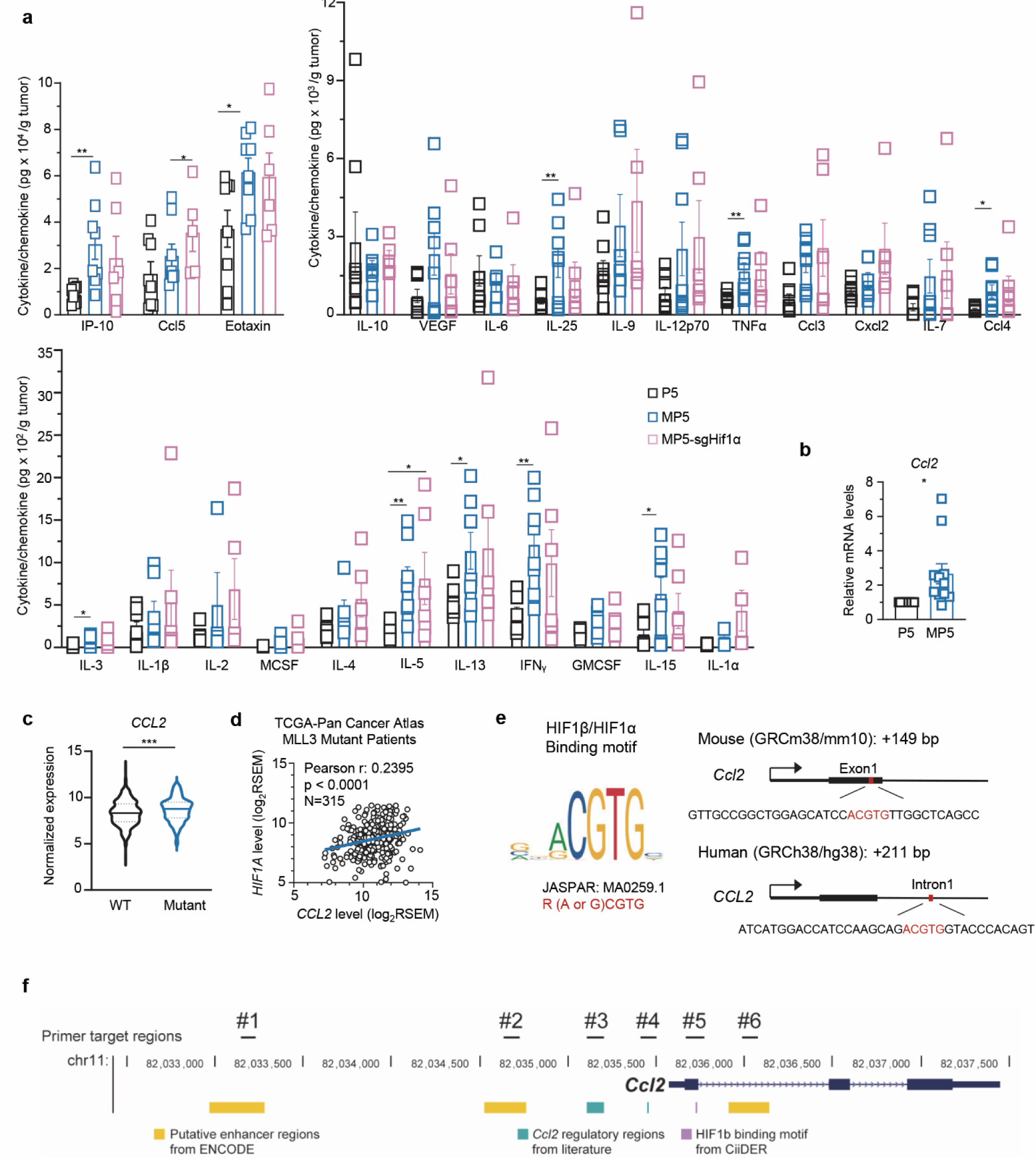
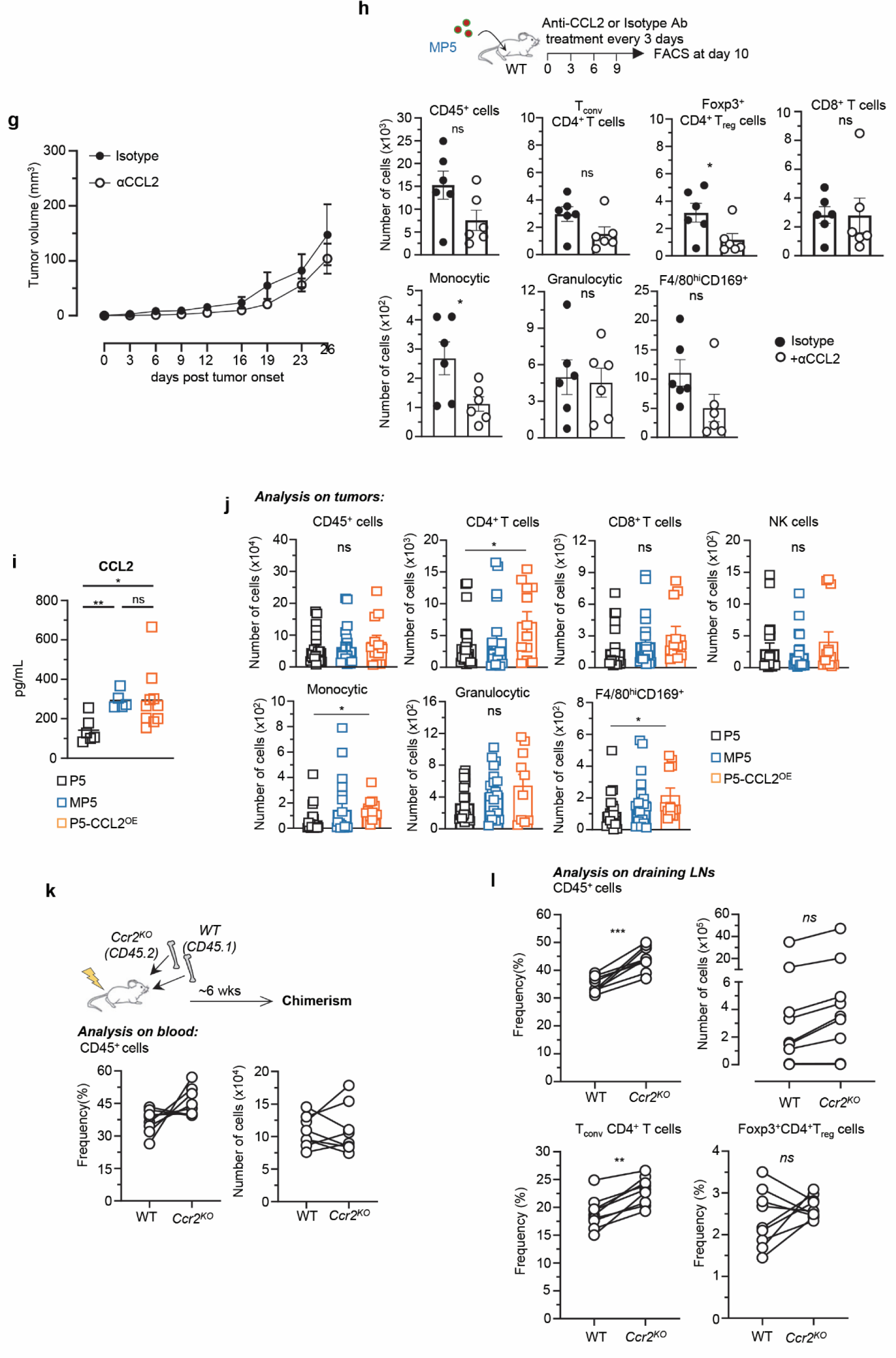
**(a)** Quantification of levels of chemokines (CCL2, CCL3, CCL4, CCL5, CXCL2, CXCL10 and Eotaxin), cytokines (IL-1α, IL-1β, IL-6, IL-12p70, IFNγ, TNFα, IL-4, IL-5, IL-13, IL-25, IL-9, IL-2, IL-7, IL-15, IL-10 and IL-3) and growth factors (GMCSF, MCSF and VEGF) in P5-sg*NT*, MP5-sg*NT* and MP5-sg*Hif1a* pre-onset nodules as depicted in Figure 4f. **(b)** *Ccl2* mRNA expression in P5 and MP5 organoid lines #1, #2, #3, #4, and #7. **(c)** *CCL2* mRNA expression in *MLL3*-WT (n=769) and *MLL3*-mutant (n=315) patients in TCGA Pan Cancer Atlas Breast Cancer database. **(d)** Correlation of *HIF1A* and *CCL2* gene expression in MLL3-mutant patients (n=315) in TCGA Pan Cancer Atlas Breast Cancer database. RSEM, RNA-seq by Expectation Maximization. **(e)** HIF1β /HIF1α binding motif (Hypoxia response element, HRE). Motif sequence is referred from JASPAR database. This motif is conserved around *CCL2* transcription start site in both human and mouse. **(f)** Putative regulatory regions and HIF binding sites at the *Ccl2* gene locus in mouse genome. Primer amplifying regions in Figure 5c are indicated by black lines. **(g)** MP5 tumor growth as depicted in Figure 5d in a pool of 2 experiments (n=10 tumors/group). **(h)** Numbers of MP5 tumor-infiltrating immune cells and indicated subsets as determined by flow cytometry analysis in anti-CCL2 versus isotype mAb-treated mice in a pool of two replicate experiments (n=6 tumors). **(i)** Quantification of levels of CCL2 in P5-control vector (n=6), MP5-control vector (n=5), and P5-CCL2^OE^ (n=10) tumors (3 mm diameter) by ELISA. **(j)** Number of indicated immune infiltrating cell subsets as determined by FACS in pre-onset tumor nodules of FVB mice transplanted with P5-control vector, MP5-control vector and P5-CCL2^OE^ MaSCs as depicted in Fig. 5f. **(k, l)** Analysis of leukocytes (CD45^+^) chimerism (frequency and number) in the blood of mixed-BM *Ccr2^-/-^*/WT chimeras, as depicted in Fig. 5g (k) and in tumor-draining LNs, including CD4^+^ T_conv_ and T_reg_ cells (l). Each line corresponds to 1 individual mouse in 1 of 2 replicate experiment. P-values were determined by unpaired t-test (a,b,c,h,i and j) or Pearson’s rank correlation (d) or Log-rank test (g) or paired t-test (l). *p< 0.05, **p< 0.01, ***p< 0.001, ns or not indicated: non-significant.

**Supplemental Figure 6, related to Figure 6.**
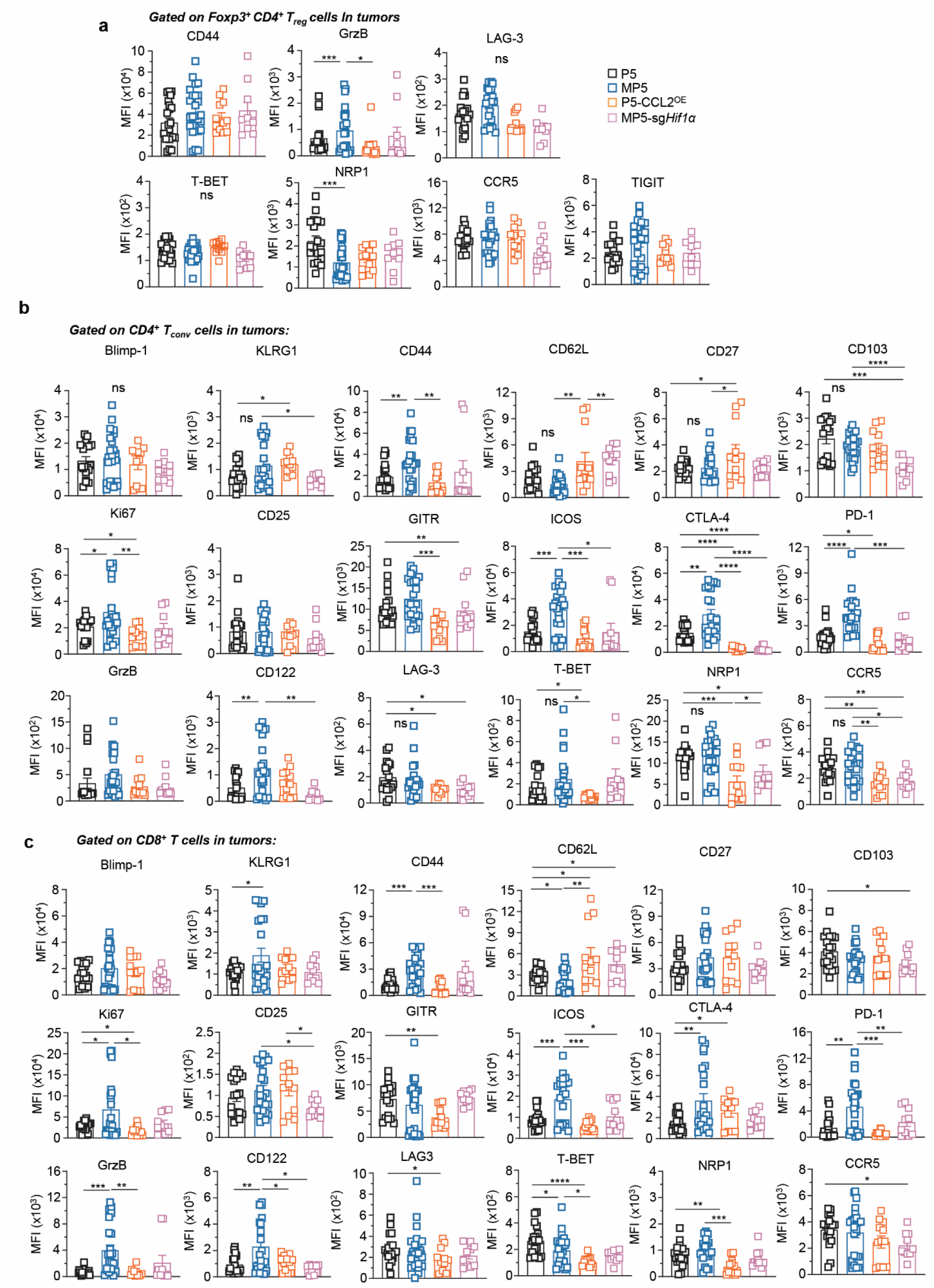
Relative average expression level of indicated markers on T_reg_ cells **(a)**, CD4^+^ T_conv_ cells **(b)** and CD8^+^ T cells **(c)** infiltrating indicated pre-onset tumors based on 22-marker flow cytometry analysis. Data are from 4 independent replicate experiments, in P5 (n=19), in MP5 (n=22), in P5-CCL2^OE^ (n=13), in MP5-sg*Hif1α* (n= 9) tumors. Each symbol (bar graphs) feature 1 tumor. P-values were determined by unpaired t-test (a-c). *p< 0.05, **p< 0.01, ***p< 0.001, **** p< 0.0001, ns or not indicated: non-significant.

**Table S1: List of all antibodies used in the study**

**Table S2: Breast tumor patient information**

**Table S3: List of all primers used in the study**

